# Microbiome signatures in *Acropora cervicornis* are associated with genotypic resistance to elevated nutrients and heat stress

**DOI:** 10.1101/2022.05.02.490297

**Authors:** Ana M. Palacio-Castro, Stephanie M. Rosales, Caroline E. Dennison, Andrew C. Baker

## Abstract

The staghorn coral, *Acropora cervicornis*, was once abundant in the Caribbean, but now is listed as critically endangered. To recover *A. cervicornis* populations, restoration efforts have focused on preserving genetic diversity and increasing coral cover. However, identifying stress-resistant corals can help to increase restoration success, by allocating genotypes to reefs where they are more likely to survive. We assessed the performance (growth, survivorship, and photochemical efficiency) and characterized the microbiome (prokaryotes) of six *A. cervicornis* genotypes that were maintained at control temperatures (~26 °C) and either ambient nutrients or elevated nutrients (elevated NH_4_, and elevated NH_4_ + PO_4_) for > 2 months. We then compared how these parameters changed when the corals were exposed to heat-stress (3-weeks at ~31.5 °C). We found that exposure to elevated nutrients reduced *A. cervicornis* performance under control temperatures and heat stress. However, there was a wide range of variation among genotypes, with three genotypes maintaining relatively higher survivorship and growth rates when exposed to nutrients alone, and nutrients followed by heat stress. Heat stress alone changed the microbial composition among genotypes more than elevated nutrients alone, but heat stress also interacted with nutrient pre-exposure to affect microbial communities. The relative abundance of *Midichloriaceae* and *Spirochaetaceae* varied by coral genotype and a high abundance of these bacterial taxa was a positive predictor of coral survivorship rate, suggesting a microbial signature that could aid in identifying resistant *A. cervicornis* genotypes. Our findings suggest there is significant variation among genotypes in the response of *A. cervicornis* to elevated nutrients and temperatures. Resistant genotypes may be identifiable via their microbiomes and prioritized for outplanting at sites characterized by high levels of nutrient pollution. Large-scale microbiome screening may help expedite targeted outplanting and could be tested and extended to facilitate the identification of genotypes with other resistance characteristics.

## Introduction

Coral cover in the Caribbean has declined in recent decades due to natural and human disturbances (Gardner et al. 2003). However, *Acropora cervicornis* populations have been particularly affected, with an estimated cover loss of up to 90% between the 1970s and 1990s (Precht et al. 2002). *A. cervicornis* is now listed as ‘threatened’ under The United States Endangered Species Act (NMFS 2006) and is the target of restoration programs that aim to preserve its genetic diversity and increase coral cover. To achieve this, coral fragments are currently propagated in nurseries and later out-planted to restoration sites (Young et al. 2012). The ultimate goal is for the fragments to grow and become sexually mature to self-sustain functional coral populations (Lirman and Schopmeyer 2016).

However, one limitation for coral restoration is that reefs continue to face local and global stressors. Water quality issues, for example, are common in reefs near metropolitan areas and can be exacerbated by coastal development, increasing agricultural runoff, and sea-level rise (McKenzie et al. 2021). Elevated nutrients can reduce coral calcification (Renegar and Riegl 2005) and increase bleaching susceptibility (Wiedenmann et al. 2013). Nutrients have been also reported to increase disease prevalence (Vega Thurber et al. 2013), and disrupt corals’ associated microbial communities, altering the cycling of nutrients among the coral host and their symbionts (Rädecker et al. 2015) and promoting the proliferation of opportunistic or pathogenic microbes (Vega Thurber et al. 2009). However, the effects of nutrient enrichment on corals are not always negative and can vary by coral species (Fox et al. 2021; Palacio-Castro et al. 2021), nutrient levels (D’Angelo et al. 2014; Dobson et al. 2021), nutrient source (Burkepile et al. 2019), exposure time (Fabricius 2005), and the ratio of nitrogen to phosphorus availability (Ferrier-Pagès et al. 2000; Rosset et al. 2017).

On a global scale, climate change is the biggest threat to coral reefs, and it is causing more frequent and severe bleaching events (Eakin et al. 2019). Sustained seawater temperatures 1-2 °C above the average local summer maximum can trigger the breakdown of the coral-algal symbiosis (Warner et al. 1999), resulting in the loss of the alga and in the paling of the coral (Glynn 1993). Because up to 90% of the coral’s carbohydrate supply comes from their algal symbionts (Muscatine and Porter 1977), bleached corals are in a physiologically and nutritionally compromised state that usually leads to coral mortality (Baker et al. 2008). Sublethal bleaching also results in lower coral growth (Goreau and Macfarlane 1990), reproductive output (Ward et al. 2002), and disease resistance (Muller et al. 2018). Heat stress also alters the structure and diversity of the prokaryotic communities associated with corals, and commonly results in more diverse communities with higher abundances of opportunistic bacteria (McDevitt-Irwin et al. 2017).

Since climate change will continue to occur for some decades, even if global emissions are immediately halted (Donner 2009), it is imperative to find mechanisms that facilitate the persistence of coral reefs. Interventions, such as identifying corals with enhanced stress resistance, can help to increase the survivorship of endangered species and coral outplants (van Oppen et al. 2017; Goergen and Gilliam 2018; NASEM 2019). This process includes assessing the performance and stress response of different genotypes, characterizing the interaction of these genotypes with the environment (phenotypic plasticity; Drury et al. 2017), and establishing the role of each holobiont member (host, endosymbiotic algae, bacteria, archaea, and viruses) in the resulting phenotypes. Genotypic differences in *A. cervicornis* are documented in disease susceptibility (Miller et al. 2019), bleaching, and bleaching recovery (Muller et al. 2018). In *A. cervicornis,* outplant success has been evaluated by tracking *Symbiodinium ‘fitti’* strains but no correlation was found between survivorship and *S. fitti* strain (O’Donnell et al. 2018). In contrast, members of the microbial community have been linked to patterns of disease and bleaching resilience (Chu and Vollmer 2016; Klinges et al. 2020).

In a companion study, we found that *A. cervicornis* is particularly sensitive to the combined effects of elevated nutrients and temperature (Palacio-Castro et al. 2021). Here we investigated genotypic variation in sensitivity by comparing the performance (growth, survivorship, and photochemical efficiency), and prokaryotic community (bacteria and archaea) of six *A. cervicornis* genotypes that are used in coral restoration in South Florida. These genotypes were studied under: (1) control temperatures and ambient nutrients, (2) control temperatures and elevated nutrients (NH_4_ and NH_4_ + PO_4_), (3) elevated temperature in corals pre-exposed to ambient nutrients, and (4) elevated temperature in corals pre-exposed to elevated nutrients. Specifically, we tested whether some *A. cervicornis* genotypes are more resistant to elevated nutrients and temperature, and if the composition of their microbial communities can explain some of the variations in *A. cervicornis* stress response.

## Methods

### Coral collection

Single-branched fragments from six *A. cervicornis* genotypes were donated by the University of Miami and Mote Marine Laboratory nurseries in summer 2017 (N=120, 8-29 fragments per genotype; Table S1). These corals were transported to the Marine Technology and Life Science Seawater (MTLSS) complex at the University of Miami Rosenstiel School in September 2017, where they were acclimated to tank conditions at 26.2 °C (±0.6 SD) for ~4 months.

### Experimental conditions

Phase 1 (days 1-78) - Control temperature and nutrient treatments: After acclimation, all fragments per genotype were evenly assigned to one of six 38-L glass aquaria which were divided between two independent tanks. The tanks acted as a water bath to maintain the target temperature at ~26 °C (26.1 °C ± 0.4 SD) and contained one aquarium (replicate) of each nutrient treatment. Fragments in the ambient treatment (ambient) were maintained under nutrient concentrations from the incoming water inflow from Biscayne Bay, FL (NH_4_ = 1.18 μM ± 0.97 SD, PO_4_ = 0.18 μM ± 0.12 SD). Corals in the NH_4_ treatment were dosed with NH_4_Cl to increase the ammonium concentration by ~10 μM over the ambient values (NH_4_ = 10.22 μM ± 2.01 SD, PO_4_ = 0.15 μM ± 0.01 SD). Corals in the NH_4_ + PO_4_ treatment were dosed with NH_4_Cl in a similar way, plus with NaH_2_PO_4_H_2_O to increase phosphate concentration by ~1 μM relative to ambient (NH_4_ = 10.38 μM ± 1.13 SD, PO_4_ = 0.73 μM ± 0.15 SD). These nutrient concentrations are in the range of nutrient levels that South Florida can experience during high nutrient periods (Lapointe et al. 2004; Caccia and Boyer 2005).

Phase 2 (days 79-90) - Ramp-up temperature and nutrient treatments: During this phase, corals were maintained in their respective nutrient treatments (ambient, NH_4_, and NH_4_ + PO_4_), while the temperature in the tanks was gradually increased from 26 °C to ~31.5 °C over 12 days (Fig. 1). This target temperature was ~1°C above the maximum monthly mean temperature in South Florida reefs (Gintert et al. 2018), which is considered the temperature threshold for the accumulation of heat stress in corals (Liu et al. 2006).

**Fig. 1:**
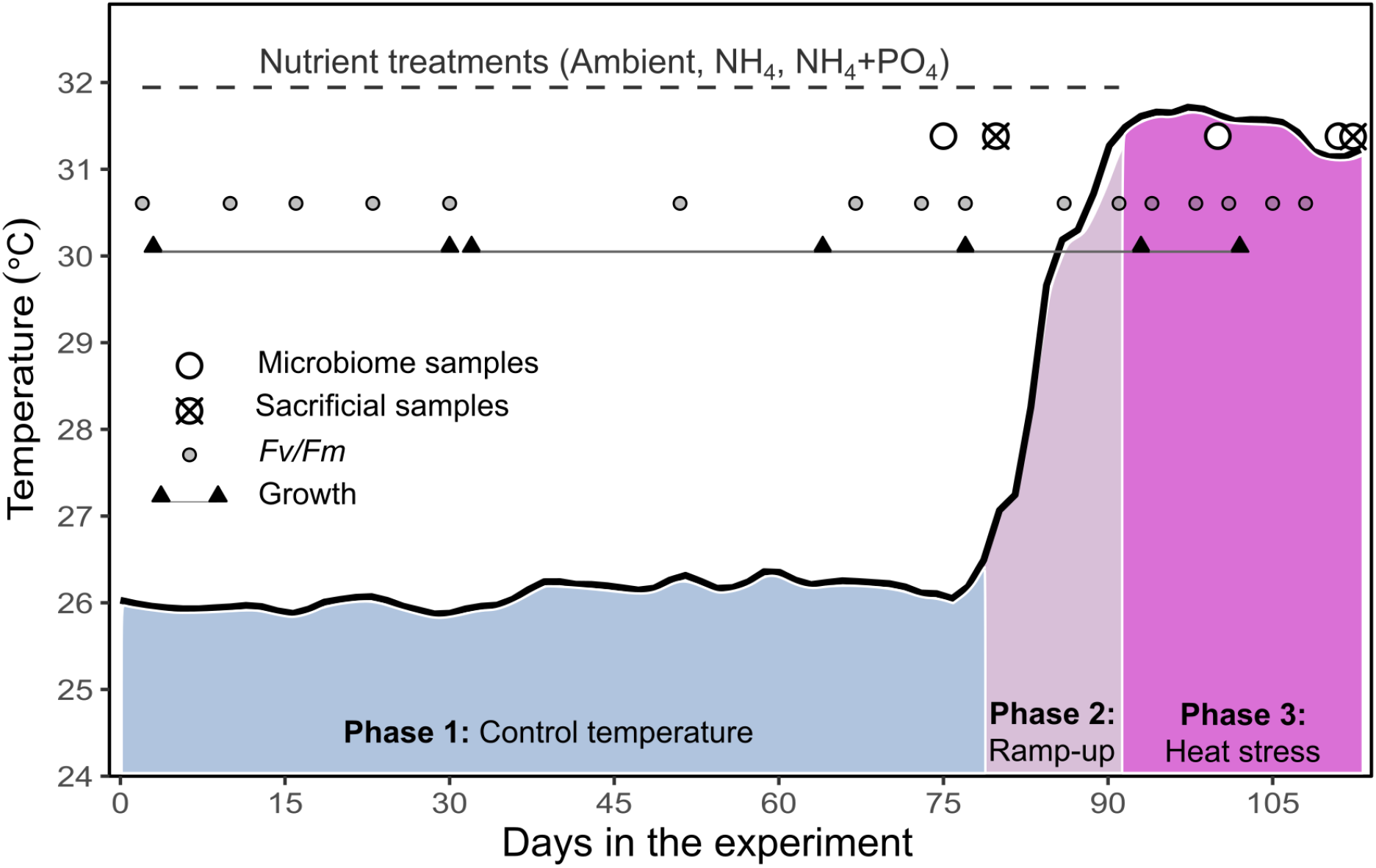
Experimental conditions and days when samples and data were collected. Black solid line shows the smoothed mean temperature. Top dashed line denotes the period of nutrient addition during phase 1 (control temperature) and phase 2 (ramp-up). Due to the onset of coral mortality in the elevated nutrient treatments, all the fragments were maintained under ambient nutrients during phase 3 (heat stress). Microbiome samples were non-sacrificial small tissue biopsies (~ 2 polyps) which allowed to repeatedly sample the same fragments over time. Sacrificial samples (N=39) consisted of the remotion from the experiment of a subset of fragments for a companion study.

Phase 3 (days 91-110) - Heat stress and ambient nutrients: In this phase, all the fragments were maintained at 31.5 °C (±0.8 SD). The initial goal was to maintain the nutrient treatments through heat stress, but given a high level of coral mortality in elevated nutrients (NH_4_, and NH_4_ + PO_4_), nutrient additions were halted on day 91 (Fig. 1).

### Coral performance

Coral survivorship, growth rates, and photochemical efficiency (*F_v_/F_m_*) were used as proxies of genotype performance under the nutrient treatments (ambient, NH_4_, and NH_4_ + PO_4_), and temperature conditions (control, ramp-up, and heat stress; Fig. 1). Fragment survivorship was monitored daily by recording when a fragment died or was sacrificed to collect samples. Buoyant weight data were collected once a month before the experiment (baseline) and monthly during phase 1. During phases 2 and 3, buoyant weight was measured approximately every two weeks. The dark-adapted photochemical efficiency of photosystem II (*F_v_/F_m_*) was recorded approximately every two weeks during phase 1, and twice a week during phases 2 and 3, using an Imaging-Pulse Amplitude Modulated Fluorometer (I-PAM, Walz, Effeltrich, Germany). Details on data collection are described in the ESM.

Coral performance data analyses were conducted in R v3.6.3 (R Development Core Team 2020). We first tested for differences among the corals maintained in all three nutrient treatments (ambient, NH_4_, and NH_4_ + PO_4_) over time, but the survivorship, growth rates, or *F_v_/F_m_* values of the fragments maintained in NH_4_ compared to NH_4_ + PO_4_ were not different. For simplicity, these two treatments were pooled and presented as an “elevated nutrients’’ treatment that was used to test for differences among the genotypes when exposed to ambient versus high nutrients (results for NH_4_ and NH_4_ + PO_4_ separated are shown in Figs. S1-S3).

Survivorship probabilities were estimated with survival 2.38 (Therneau 2015) and survminer 0.4.6 (Kassambara 2018) packages for R. Log-rank tests were used to test the additive effects of nutrients treatments (ambient versus elevated nutrients) and genotypes on corals’ survivorship. Additionally, a Cox proportional model was used to estimate the relative hazard ratio (HR) of the genotypes when exposed to elevated nutrients and heat stress. The HR is calculated as the ratio of the total number of observed to expected deaths between two genotypes and the resulting ratio represents the number of times that the risk of death is higher in one genotype compared to another. Thirty-nine fragments were sacrificed during the experiment (28 at the end of phase 1, and 11 at the end of phase 3; Fig. 1) for a companion study (Palacio-Castro et al. 2021), and their removal was recorded as “censored” events to adjust the survivorship curves. These events account for the incomplete information about the survivorship outcome of the removed fragments since it is unknown if they would die or survive if left longer in the experiment. Information about the “censored” fragments is then incorporated into the model until they are removed, but they are “censored” after that day (i.e. not considered as part of the sample groups).

Differences among the growth rates and *F_v_/F_m_* in the treatments and genotypes were evaluated with linear mixed models. The models were run with the lme4 package v1.1-17 (Bates et al. 2015) and pairwise comparisons among significant effects with emmeans (Lenth 2018) using an alpha value of 0.05 for the Tukey’s HSD contrasts. Each model included genotype, day, and nutrient treatment (ambient versus elevated nutrients) as fixed factors, as well as coral fragment and replicate tank as random effects (Tables S3 and S6 show model outputs, and S4, S5 and S7 show the post-hoc tests). Performance data and code for data analyses are available at Zenodo (https://doi.org/10.5281/zenodo.5041424).

### Prokaryotic alpha and beta diversity

A subset of fragments was sampled at the end of phase 1 (day 75, n=85), and during phase 3 (days 100 and 111, n=52 and n=42, respectively) to characterize prokaryotic communities in each genotype and treatment (Table S2). The selected fragments in phase 1 were re-sampled in phase 3. However, the number of samples decreased through time because some of the fragments died prior to re-sampling. In these samples, the 16S rRNA gene V4 was amplified and sequenced using previously published primers (Apprill et al. 2015). Details on the prokaryotic library generation and bioinformatics processing can be found in the ESM. For both alpha (the variation of prokaryotic members) and beta diversity (prokaryotic community composition), the data was parsed by multiple categories: (1) differences among genotypes were compared on ambient fragments on day 75 [phase 1] since these corals did not undergo nutrient or temperature stress, (2) nutrient treatments at control temperature were compared in ambient, NH_4_, and NH_4_+PO_4_ fragments at day 75 [phase 1], (3) nutrient treatments at elevated temperatures were compared in ambient, NH_4_, and NH_4_+PO_4_ fragments at days 100 and 111 [phase 3], and (4) each nutrient treatment was compared across days (days 75, 100, and 111). For across days data, only genotypes that survived through the end of the heat stress (*G48, G62* in NH_4_, and *G48, G62, G31* in NH_4_+PO_4_) were used. In ambient corals, all genotypes survived and were evaluated through time. Both alpha and beta-diversity analyses were considered significant if alpha was <0.05.

For alpha diversity, the breakaway plugin on Qiime2-2019.7 was used to calculate the prokaryotic richness in each category without normalizing by sequencing depth (rarefraction; Willis et al. 2017) since this can introduce biases (Willis 2019). Richness values were then square root-transformed and tested with linear mixed models in the lme4 package (v1.1.21) for R. Pairwise Tukey’s HSD comparisons were evaluated with emmeans (v1.4.3.1).

For beta-diversity analysis, the count table was filtered to remove ASVs present in ≤ 4 fragments. The ASV count table was normalized to centered log-ratio (CLR) values with the R package microbiome v1.4.2 (Lahti et al. 2017), which is recommended given the compositional structure of the amplicon sequence data (Gloor et al. 2017). Transformed data were used to compute a dissimilarity matrix based on Euclidean distance using the function vegdist (Vegan v2.5.6 package; Dixon 2003). To identify differences between-group beta-diversity, the dissimilarity matrix was tested with the function Adonis (PERMANOVA; Vegan v2.5.6) with 999 permutations and the option strata to control for genotype effect. Pairwise comparisons were conducted with the package pairwiseAdonis v0.0.1 with a Bonferroni correction (Martinez Arbizu 2017). The dispersion of the dissimilarity matrix was calculated using the function betadisper (Vegan v2.5.6) and samples were tested for within-group beta-diversity differences with an ANOVA and a post hoc test (Tukey’s HSD). In addition, the entire dataset was evaluated to investigate which factor and interaction (i.e., nutrients, temperature, or genotype) explained the majority of the variance.

### Prokaryotic core microbiome

The baseline (ambient corals at day 75; n = 30) core microbiome was estimated for each genotype to determine taxa that may be important for genotype health. The function ‘core’ from the R microbiome package v1.0.8 was used to determine which ASVs were present in at least 99% of fragments (at any abundance) per genotype. The number of core ASVs per genotype and the intersection among the genotypes were assessed.

### Prokaryotic differential abundance

To evaluate differentially abundant prokaryotic taxa the filtered count table (ASVs present in ≥ 4 fragments) was analyzed with the R package ANCOM2. Results were considered significant if alpha was <0.05 with a detection cut-off of 0.90 (Mandal et al. 2015). The data were first tested for changes due to nutrient treatment at day 75 with a genotype interaction. Then, differences in nutrient treatments across days 75, 100, and 111 were tested, but not for *G08, G07*, or *G50* due to their early mortality. Individual fragments were included as a random effect for all tests.

Finally, we tested differentially abundant taxa among coral genotypes against their associated survivorship outcomes under combined nutrient and heat stress (i.e., Do baseline prokaryotes correlate with coral mortality?). To do this, baseline microbiomes (ambient corals at day 75; n = 30) were evaluated with ANCOM2. The survivorship rates were generated from day 110 after corals were exposed to elevated nutrients and heat stress (i.e., NH_4_ and NH_4_+PO_4_ survivorship pooled at day 110). Survivorship rates were also categorized into high and low survivorship and tested for differential abundance. The significant taxa were then correlated to the same survivorship rates used in the ANCOM2 analysis using the lm function in R. The bioinformatic scripts for the microbiome analysis are publicly available (https://github.com/srosales712/AcNutrients).

## Results

### Coral survivorship

There was no mortality in any of the *A. cervicornis* genotypes when they were maintained in ambient nutrients. Although mortality was slightly higher among corals exposed to NH_4_ compared to NH_4_+PO_4_, there were no significant differences between the survivorship probabilities in these two treatments (log-rank p > 0.5, Fig. S1) and they were pooled as “elevated nutrients.” Survivorship probabilities in elevated nutrients were lower for genotypes *G50, G07*, and *G08* compared to genotypes *G31, G62*, and *G48* (log-rank *p*<0.0001; Fig. 2a). Under elevated nutrients, *G50* and *G07* first experienced mortality at control temperature (days 65 and 71, respectively), followed by *G08* after one day in heat stress (day 92). *G31* first suffered mortality after 1 week in heat (day 96), followed by *G62* (day 103) and *G48* (day 106) after 2 weeks in heat. The Cox-hazard ratios (HR) indicated that when the corals were exposed to elevated nutrients and heat stress the risk of death was 3 times higher for genotype *G31*, and 64-136 times higher for *G50, G07*, and *G08* compared to genotypes *G62* and *G48*, which had the lowest risk of death (Fig. 2b).

**Fig. 2:**
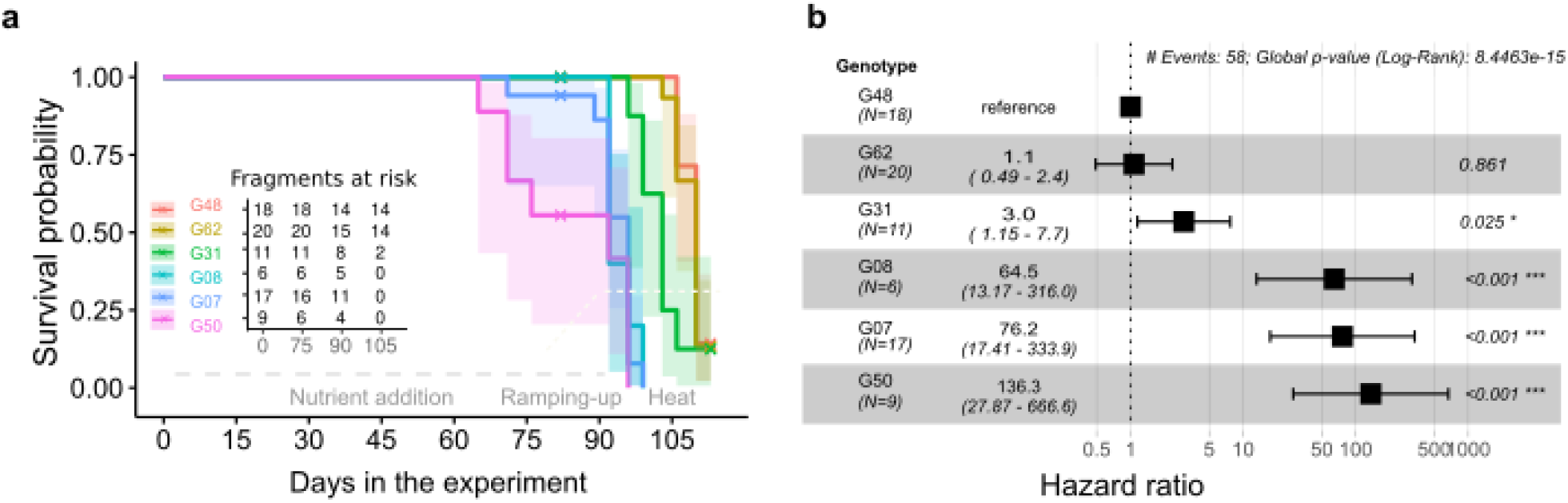
Survival probability and relative risk of death in six *Acropora cervicornis* genotypes exposed to elevated nutrients and subsequent heat stress. **a**. Survival probability of individual genotypes exposed to elevated nutrients (NH_4_ and NH_4_+PO_4_ treatments combined) at control and ramp-up temperatures (days 1-90), and during a combination of heat stress and pre-exposure to elevated nutrients (days 91-113). The “x” symbols show the days with “censored events” when fragments were removed from the experiment and therefore are not considered as part of the sample groups after that day. The “fragments at risk” table shows the number of fragments that remained in the experiment on any specific day (initial number of fragments minus fragments that died or were removed to collect whole-tissue samples). **b**. Combined effect of pre-exposure to elevated nutrients and heat stress on the hazard ratio of the risk of death of different *A. cervicornis* genotypes (x-axis). Values are relative to genotype *G48*, which had the lowest risk of death overall.

### Overall growth rate

At control temperatures, overall growth rates in the ambient treatment ranged from 3.36-3.61 mg g^-1^d^-1^. However, growth rates progressively declined under elevated nutrients compared to ambient (Fig. 3a). During the first month of nutrient exposure, growth rates were reduced by 28% in NH_4_ (*p*<0.0001), and by 16% in NH_4_+PO_4_ (Tukey’s HSD *p* > 0.05) with respect to ambient nutrient conditions. By the second month, growth was ~52-53% lower in both elevated nutrient treatments (~1.6 mg g^-1^d^-1^ ±0.2; Tukey’s HSD *p*<0.0001) with respect to ambient (Fig. 3a, Table S4).

**Fig. 3:**
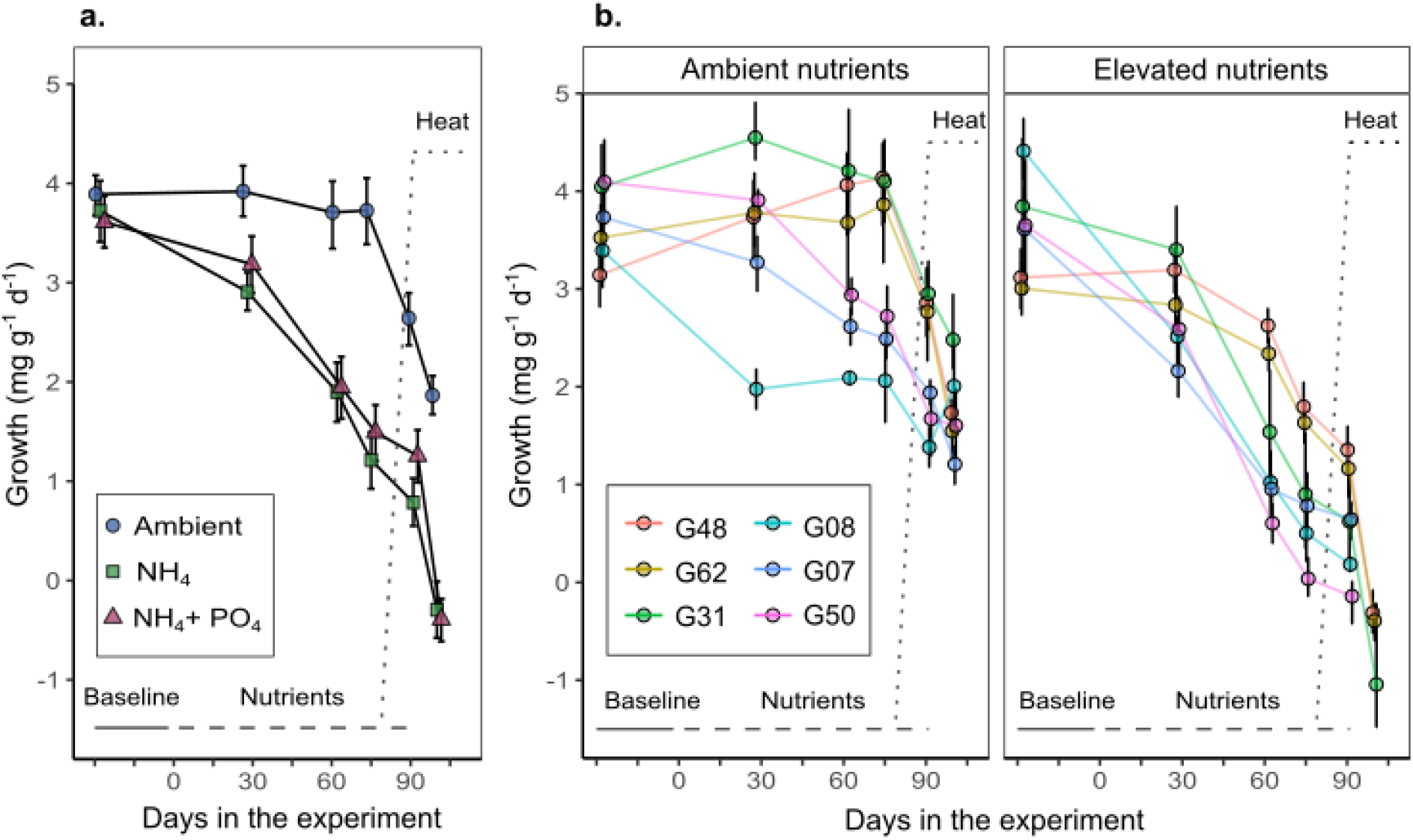
Mean growth rate of *A. cervicornis* (mg g^-1^ d^-1^ ± 95% CI) before nutrient addition (baseline), during phase 1 (control temperature), phase 2 (ramp-up), and phase 3 (heat stress)**. a.** Overall growth rates (all genotypes pooled by nutrient treatment). **b.** Genotype-specific growth rates by nutrient treatment. Elevated nutrients panel includes the data from NH_4_ and NH_4_+PO_4_. The gray solid line demarcates the baseline measurement, the dashed line the nutrient addition period, and the dotted line the ramp-up and heat stress periods.

Heat stress reduced growth rates in both ambient and elevated nutrients (Fig. 3a). Corals in ambient nutrients experienced a 30% growth decline during ramp-up (phase 2, Tukey’s HSD *p*<0.0001), and a 52% decline after a week in heat stress with respect to their last phase 1 values (*p*<0.0001). By the end of heat stress, the mean growth rate in ambient nutrients was 1.61 mg g^-1^d^-1^ ±0.2. Heat stress further reduced growth rates in corals pre-exposed to elevated nutrients. After a week of heat stress, NH_4_ and NH_4_+PO_4_ fragments reached negative growth values (dissolution) (−0.69 and −0.79 mg g^-1^d^-1^ ±0.26, respectively; Fig. 3a, Table S4).

### Genotype growth rates

Differences in growth rate between corals in the two elevated nutrient treatments were non-significant at all time points (Tukey’s HSD *p* > 0.05; Figs. 3a, S2). Based on this, NH_4_ and NH_4_+PO_4_ data were pooled under “elevated nutrients”. Growth rates varied among genotypes, and there was an interaction between genotype, nutrients, and temperature (Fig. 3b). Baseline growth (before nutrient addition) for *G48* and *G62* (3.13 - 3.26 mg g^-1^d^-1^) was lower than for other genotypes (3.88 - 4.22 mg g^-1^d^-1^; Tukey’s HSD *p*<0.05), but this changed during the experiment. Under ambient nutrients, *G48* experienced up to a 32% increase in growth during phase 1, while *G50*, *G07*, and *G08* experienced a 33-39% reduction in growth (Fig. 3b). By the end of phase 1, the hierarchical ranking of genotype growth rate in ambient was inverted, with respect to baseline (pre-experiment) data, and *G48*, *G31*, and *G62* had higher growth rates (3.86 - 4.14 mg g^-1^d^-1^) than *G50*, *G07*, and *G08* (2.06-2.72 mg g^-1^d^-1^; Tukey’s HSD *p*<0.05; Fig. 3b, Table S5). During phase 3 (heat stress: days 91-100), *G31* in ambient was the only genotype that maintained growth rates > 2 mg g^-1^d^-1^ (~2.5 mg g^-1^d^-1^ Fig. 3b).

When exposed to elevated nutrients, *G48* and *G62* had the lowest reductions in growth rate with respect to their values in the ambient treatment. By the end of phase 1, these genotypes maintained growth rates >1.6 mg g^-1^d^-1^ (~57-58% reduction with respect to ambient values at the same time). The remaining genotypes had growth rates <1.0 mg g^-1^d^-1^ at this same time (~68-105% reduction with respect to ambient values; Fig 3b, S2). Under heat stress (phase 3), only *G48*, *G62,* and *G31* had surviving fragments in the elevated nutrient treatments and all were exhibiting negative growth values (dissolution), with *G31* showing the strongest growth rate decline (Fig. 3b).

### Photochemical efficiency (*F_v_/F_m_*)

*F_v_/F_m_* values were affected by genotype, nutrient treatment, temperature, and their interaction

(Fig. 4). In the ambient treatment, *F_v_/F_m_* was lower for *G07* compared to the rest of the genotypes (Tukey’s HSD *p*<0.05). This pattern was maintained during phase 1 (−9% to −14% *F_v_/F_m_* in *G07* compared to other genotypes), phase 2 (−10 to −15%) and phase 3 (−8% to −16%). After three weeks of heat stress, *G07* had the lowest *F_v_/F_m_* in ambient nutrients (0.35±0.01), followed by *G31* (0.39±0.01), while other genotypes maintained values higher than 0.4 (Fig. 4).

**Fig. 4:**
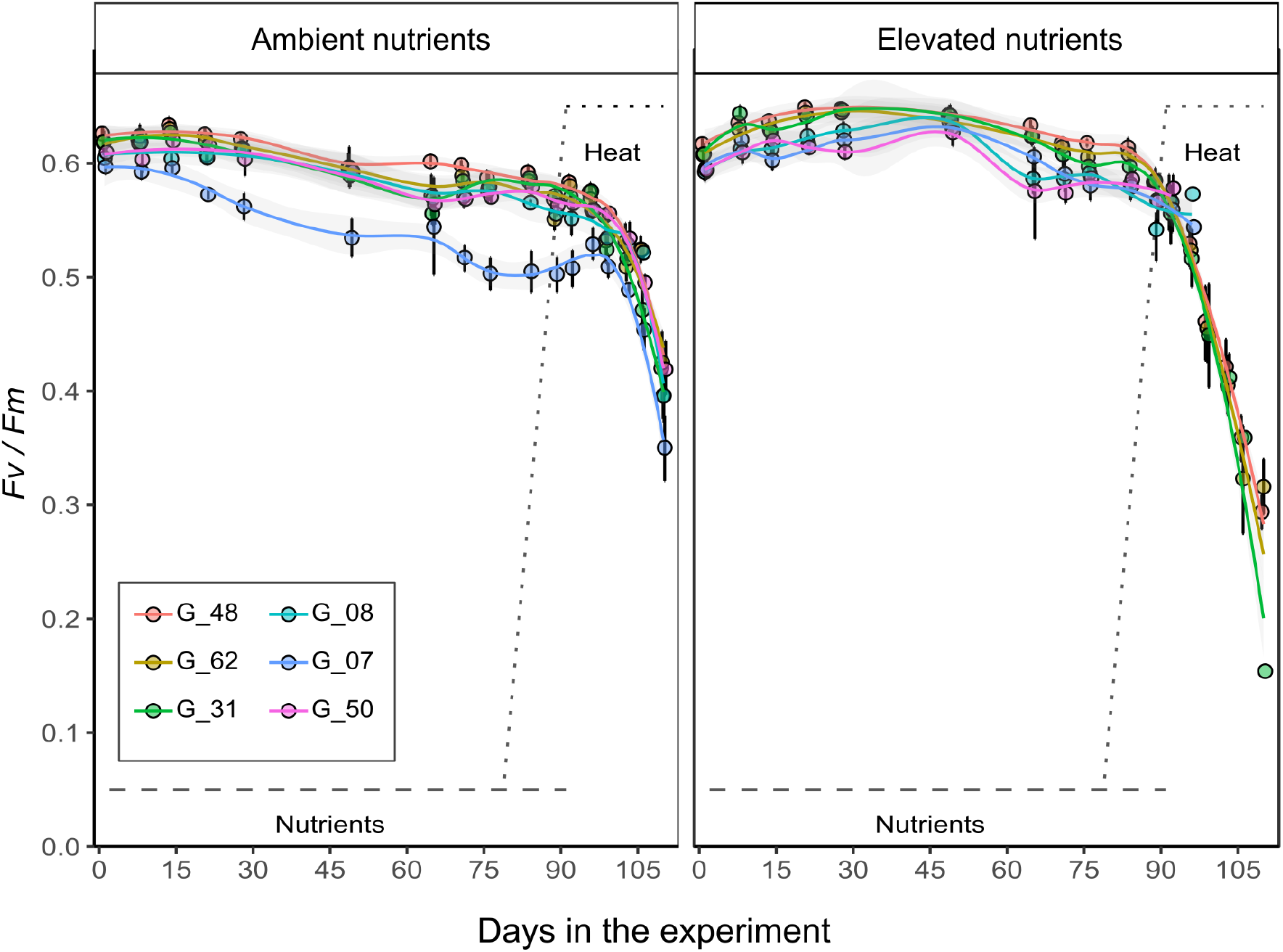
Photochemical efficiency (mean *F_v_/F_m_* ± 95% CI) of *A. cervicornis* genotypes under nutrient treatments and temperature phases. Elevated nutrient panel includes the data from NH_4_ and NH_4_+PO_4_. The gray dashed line demarcates the nutrient addition period, and the dotted line the ramp-up and heat stress periods.

During phase 1, all genotypes had a slight increase in *F_v_/F_m_* under elevated nutrients compared to ambient (2-10%), but *G07* showed the highest increase (7-18%), to the point that its *F_v_/F_m_* was not different from the rest of the genotypes in elevated nutrients (Tukey’s HSD *p* > 0.05). The positive effect of nutrients on *F_v_/F_m_* was reversed during heat stress (Figs. 4, S3). Genotypes *G50*, *G07*, and *G08* in elevated nutrients died during the first week in heat stress before *F_v_/F_m_* was assessed. In the remaining genotypes, *F_v_/F_m_* was lower in the corals pre-exposed to elevated nutrients compared to ambient nutrients. After a week in heat, *G48*, *G62*, and *G31* in elevated nutrients had 14-17% lower *F_v_/F_m_*, compared to ambient (Tukey’s HSD *p*<0.05). After three weeks in heat, these genotypes had 32%, 26%, and 62% lower *F_v_/F_m_* respectively, in nutrients compared to ambient (Fig. 4).

### Prokaryotic differences among genotypes

A total of 666 ASVs remained after filtering. Overall, corals showed a median frequency of 7,171 ASVs among the 180 samples. A comparison of the prokaryotic structures among genotypes (control temperature and ambient nutrients, n = 30) showed that *A. cervicornis* microbial communities were mainly dominated by the family *Midichloriaceae* (mean relative abundance [RA] range = 50.1-94.8%; Fig. 5a) and the core microbiome analysis showed that *Midichloriaceae* was also the only core microbe across genotypes (Fig. S4). However, genotype *G07* had the lowest RA of *Midichloriaceae* (1.4%±1.2%) and showed higher RAs of the families *Endozoicomonadaceae* (11.4%±7.1%) and *Paenibacillaceae* (9.2%±8.3%; Fig. 5a). A differential abundance analysis among genotypes showed that two ASVs from the families *Spirochaetaceae* (ASV 825; phylum Spirochaetes and genus *Spirochaeta)* and *Midichloriaceae* (ASV 209; phylum Proteobacteria and genus MD3-55) distinguished genotypes (Fig. 5b). Pairwise comparisons of these corals also showed that *G07* was the only genotype that was different from all other genotypes in alpha-diversity (padj <0.05; Fig. 5c) and dispersion (permutest; padj <0.05; Fig. 5d). Beta-diversity between genotypes was also significant (PERMANOVA; *p*=0.006), but a pairwise comparison was only significant between *G07* and *G62* (PERMANOVA; padj <0.05). *G62* and *G31* had only one core member (*Midichloriaceae*). *G48* and *G50* had three core members, but only one unique ASV among genotypes *(Spirochaeta* and *Alteromonadaceae,* respectively). *G07* had six core members and three of them were unique to *G07 (Microscillaceae* [ASV870], Acidobacteriia [ASV2724], and uncharacterized bacteria [ASV126]). *G08* had the highest number of core members (n=15), but this may have been partly driven by the low fragment number (n=2, Fig. S4). A correlation analysis between differentially abundant taxa among the genotypes and survivorship of the genotypes under nutrient and heat stress showed that both *Midichloriaceae* (*p*<0.001; R^2^=0.3), and *Spirochaetaceae* (*p*<0.0001; R^2^=0.4), had a positive correlation between their RA and survivorship (Fig. 6).

**Fig 5.**
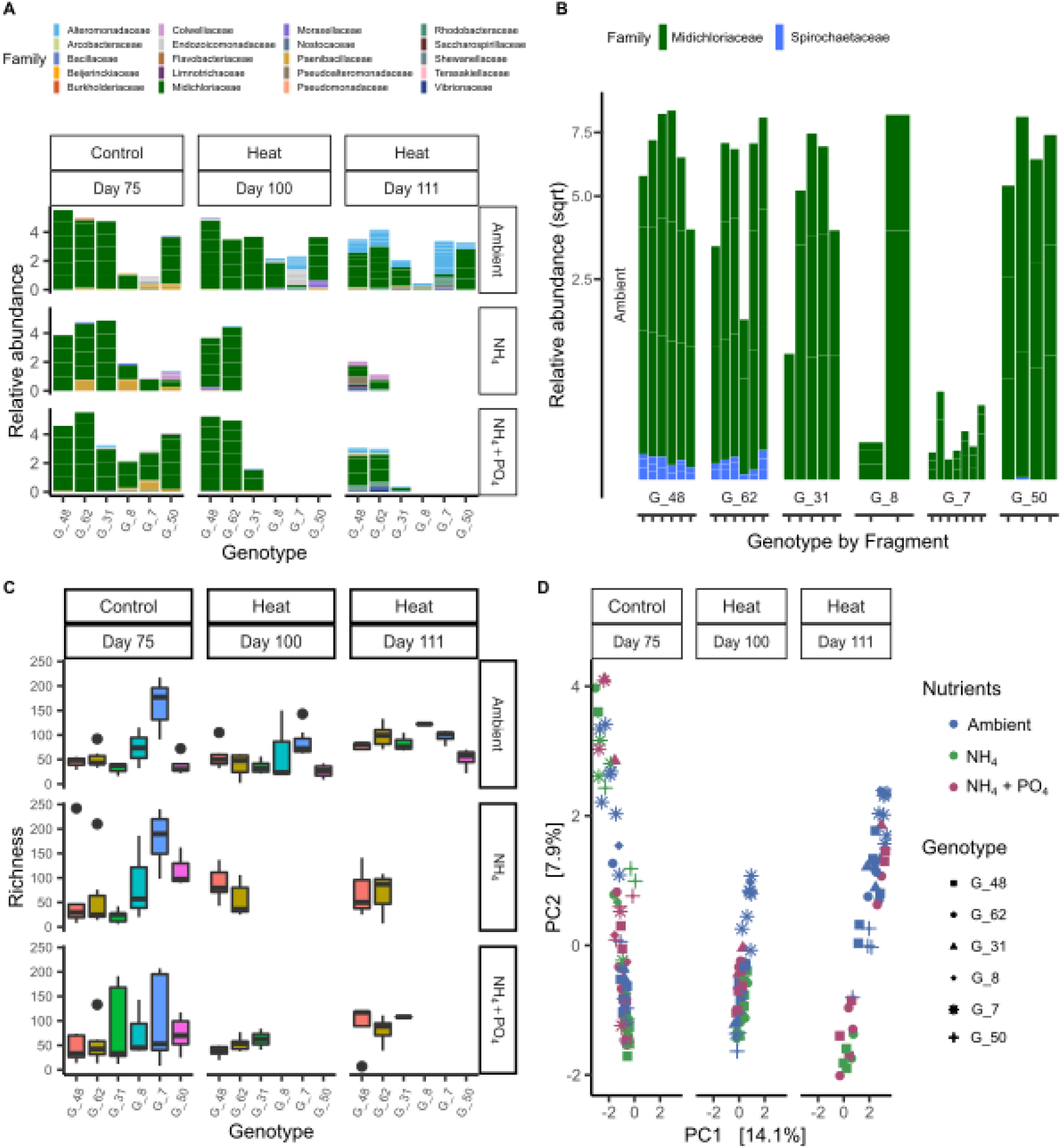
Prokaryotic communities of different *A. cervicornis* genotypes under nutrient treatments and temperature phases. (a) Relative abundance (RA; >0.05%) of the prokaryotic community of the entire dataset, (b) the RA of the two significant bacteria among the six genotypes, (c) prokaryotic richness, and (d) prokaryotic beta-diversity (centered log-ratio [CLR] transformed values on a principal component analysis [PCA] with a Euclidean distance). (a) and (c) are parsed by nutrient treatments (ambient, NH_4_, and NH_4_+ PO_4_) and heat stress (Control and Heat) across the three time points (days 75, 100, and 111). (d) is parsed by heat stress and days, the colors denote two temperature regimes and shapes denote the six genotypes and are ordered by survivorship rates.

**Fig 6.**
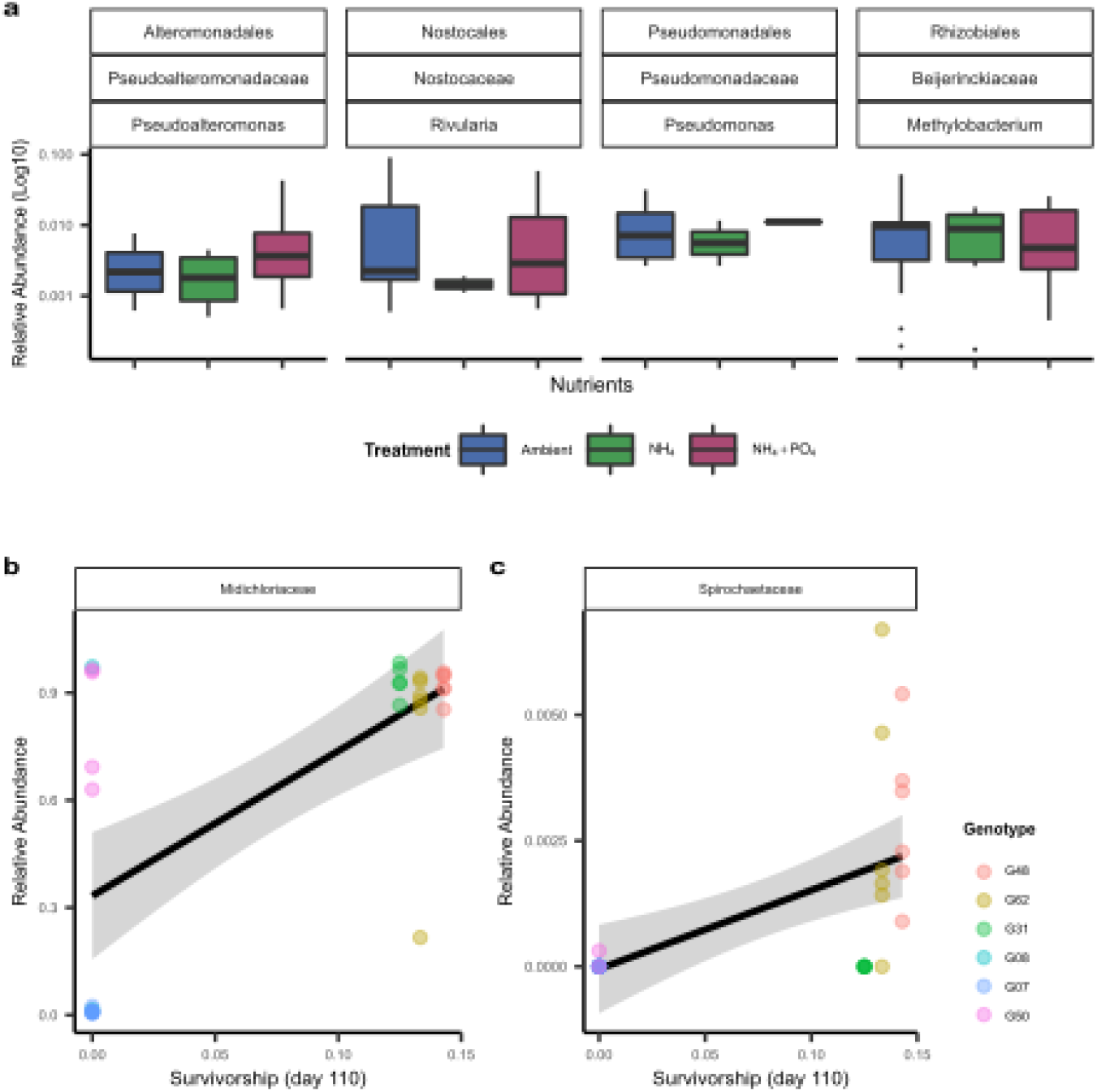
Four amplicon sequence variants (ASVs) were significantly different among nutrient treatments (phase 1; day 75). (A) Relative abundance (y-axis) of four significant taxa during nutrient treatments (x-axis; ambient, NH_4_, and NH_4_+ PO_4_) - represented by colors. The data is parsed by the ASVs corresponding order, family, and genus. Shapes denote the six genotypes and are ordered by survivorship rates on the key. A correlation analysis of (b) *Midichloriaceae* (ASV 2095; R^2^ =0.3) and (c) *Spirochaetaceae* (ASV 825; R^2^ =0.4) against the final survivorship rates (day 110) of the experiment. The colors denote the six genotypes, which are ordered by survivorship rates on the key.

### Prokaryotic beta and alpha diversity by nutrient treatment and days

Days in the experiment (temperature conditions) had the largest effect on prokaryotic composition (*p*=0.001; R^2^=0.10), followed by genotype (*p*=0.001; R^2^=0.05) and nutrient treatment (*p*=0.001; R^2^=0.03). The interaction between nutrients and genotype was also significant (*p*=0.001; R^2^=0.06). Prokaryotic beta-diversity patterns for nutrient treatments in phase 1 (day 75; n = 85) showed that nutrients alone did not change dispersion of microbial communities, but there was a group difference in beta-diversity among ambient, NH_4_, and NH_4_+PO_4_ (PERMANOVA; *p* = 0.001; R^2^ = 0.03; Fig. 5d). However, subsequent pairwise comparisons were not significant. Similarly, after nine days in heat stress (phase 3; day 100; n=38), there were no significant differences in dispersion among ambient, NH_4_, and NH_4_+PO_4_, but there was a grouping among nutrient treatments (PERMANOVA; *p*=0.001; R^2^ = 0.11) with comparisons between NH_4_+PO_4_ vs. ambient, and NH_4_+PO_4_ vs. NH_4_ being significant (padj <0.01; Fig. 5d). Unlike previous days, corals exposed to heat stress for three weeks (phase 3; day 111; n = 30) showed dispersed prokaryotic communities (permutest; *p*=0.008) and pairwise comparisons of ambient vs. NH_4_+PO_4_ was significant (permutest; padj=0.004) with NH_4_+PO_4_ showing a more dispersed community. Nutrient treatments comparisons at day 111 (phase 3) were also significant (PERMANOVA; *p*=0.001; R^2^=0.2; Fig. 6). Pairwise comparisons were significant between ambient vs. NH_4_ and ambient vs. NH_4_+PO_4_ (PERMANOVA; padj <0.03; Fig. 5d).

Beta-diversity through days for each nutrient treatment was examined independently. For all ambient corals (n = 84; days 75, 100, and 111) and NH_4_ corals (n = 26; days 75, 100, and 111), dispersion did not change across days, but beta-diversity grouping was different among days (PERMANOVA; *p*=0.001; R^2^ = 0.2; Fig. 5d). A PERMANOVA pairwise comparison was also significant for each comparison (PERMANOVA; padj <0.003). For NH_4_+PO_4_ corals (n = 41; days 75, 100, and 111), both dispersion (permutest;*p*=0.003) and grouping were different (PERMANOVA; *p*=0.001; R^2^=0.2; Fig. 5d). Pairwise comparisons showed that for NH_4_+PO_4_ corals, day 111 had higher dispersion than days 75 and 100 (permutest; padj <0.01; Fig. 5d). A PERMANOVA pairwise comparison was significant between each comparison of the three days (PERMANOVA; padj <0.01). For prokaryotic richness, a regression to nutrients showed no significant results. In ambient corals that were subsequently heat stressed, significant differences in richness were found between day 75 and day 100 (padj <0.05) and days 100 and 111 (padj <0.05) with an increase in temperature increasing richness (Fig. 5a and c).

### Prokaryotic differential abundance by nutrient treatment and days

Differential abundance analysis yielded four significant ASVs (8818, 989, 2609, and 3479) among ambient, NH_4_, and NH_4_+PO_4_ at control temperature (phase 1; day 75), but some of these differences were driven by specific genotypes (Fig. 6a). *G07* had particularly low abundances of *Midichloriaceae* (ASV 2095; 1.0%±0.7%) at control temperature (phase 1; day 75), but increased in NH_4_ (20.9%±4%) and NH_4_+PO_4_ (38.5±52.0) treatments.

Differential abundance models across days showed more differentially abundant taxa than nutrient treatments at day 75. In ambient corals across days, there were a total of 40 ASVs, followed by 14 ASVs in NH_4_+PO_4_ and 12 ASVs in NH_4_ that were differentially abundant (Fig. S5). All comparisons showed a RA increase in *Alteromonadaceae, Vibrionaceae,* and *Rhodobacteraceae* and declines in *Midichloriaceae* with days in the experiment (heat stress).

## Discussion

The rapid population decline of *A. cervicornis* has resulted in the loss of its ecological functions leading to it being a primary focus for coral restoration programs. To maximize restoration efforts, particular genotypes can be selected to re-populate certain regions based on local stressors. Here we show that there is extensive variation in the response of six *A. cervicornis* genotypes to elevated nutrients, temperature, and their combined effects, with significant differences among genotypes in their survivorship, growth rates, and photochemical efficiency. We also found genotypic differences in the prokaryotic community and used these data to explain some of the variation in the genotypic response to these stressors.

### Response to elevated nutrients

Dissolved inorganic nutrients have variable impacts on different coral species (Fox et al. 2021; Palacio-Castro et al. 2021). In this study, elevated nutrients (NH_4_ and NH_4_ +PO_4_) increased the photochemical efficiency of the algal symbionts in *A. cervicornis* but had detrimental effects on coral growth and survivorship. The mechanisms by which elevated nutrients can interfere with corals’ health are not well understood yet. However, it has been suggested that higher nitrogen availability can shift the symbiosis from a mutualistic to a parasitic relationship (Wooldridge et al. 2017; Baker et al. 2018). Higher nitrogen concentrations can benefit the algal symbionts by boosting protein synthesis and cell division (Hoegh-Guldberg 1994), leading to lower translocation of fixed carbon from the algae to the coral host and to higher symbiont cell densities (Falkowski et al. 1993; Marubini and Davies 1996). Lower calcification rates and survivorship under elevated nitrogen could then be associated with either a reduced availability of photosynthetic products for the coral (Dubinsky et al. 1990; Jiang et al. 2014) or increased competition for inorganic carbon between the coral and the algae for calcification and photosynthesis, respectively (Marubini and Davies 1996; Marubini and Thake 1999; Silbiger et al. 2018).

The combination of PO_4_ and NH_4_ did not elicit different responses in *A. cervicornis* performance compared to NH_4_ alone, suggesting that increased algal growth under elevated NH_4_ was the main driver of the physiological changes observed in both treatments. Studies that tested the effects of elevated nitrogen and phosphorus, both individually and combined, suggest that calcification could be impaired through different mechanisms under different nutrient ratios (e.g., Simkiss 1964; Ferrier-Pagès et al. 2000). In the absence of elevated nitrogen, phosphorus can act as a crystal poison directly suppressing calcification (Simkiss 1964). However, under elevated nitrogen or nitrogen and phosphorus together, both nutrients can be uptaken by the algal symbionts with the subsequent decrease in translocation of photosynthetic products to the coral (Simkiss 1964; Ferrier-Pagès et al. 2000).

Coral performance may also be shaped by the composition of the associated prokaryotic communities and their response to elevated nutrients (Shaver et al. 2017). We found that *A. cervicornis* microbiomes were marginally affected by nutrient treatment (Fig. 6a). When all genotypes were evaluated together, only four taxa were differentiated in nutrient treatments, but these taxa varied in abundance across genotypes. Similar small-to-moderate changes in corals’ prokaryotic communities have been reported for short-term (3 weeks; Maher et al. 2019) and long-term (3 years; Zaneveld et al. 2016) exposure to elevated nutrients. It is unclear why nutrients alone do not cause widespread changes in the corals’ microbial community composition, but specific genotypes show deviations from this pattern and this may be due to genotypes harboring specific microbial communities (Glasl et al. 2019). In our study, a genotype that hosted low abundances of *Midichloriaceae* in ambient nutrients (RA<2% in *G07* compared to 50.1-94.8% in other genotypes), increased *Midichloriaceae* relative abundance in elevated nutrients (mean RA=30.7%±45; Fig. 5a). This pattern has been described in other *A. cervicornis* genotypes with low baseline *Midichloriaceae* abundances (RA<12%) exposed to elevated nutrients (Shaver et al. 2017), suggesting that *A.cervicornis* have specific microbiome genotype associations (Glasl et al. 2019) that could impact their response to stressors like nutrients.

### Response to heat stress and its interaction with elevated nutrients

Heat stress alone reduced growth rates and *F_v_/F_m_* (Figs. 3, 4) but did not cause coral mortality (Fig. 2a). We did not find particularly heat-sensitive genotypes across all physiological metrics. However, heat stress alone had larger impacts on the microbial community than elevated nutrients alone, increasing the abundance of opportunistic taxa across genotypes (i.e., *Alteromonadaceae*, *Vibrionaceae*, and *Rhodobacteraceae*), a common pattern in corals exposed to elevated temperatures (McDevitt-Irwin et al. 2017). With an increase in opportunistic taxa, there was a relative decline in the dominant taxon, *Midichloriaceae*. This pattern has been described in bleached *A. cervicornis* where a reduction of *Midichloriaceae* may lead to the growth of other opportunistic taxa (Klinges et al. 2020).

The combination of heat stress and pre-exposure to elevated nutrients produced more detrimental effects on *A. cervicornis* performance than either stressor alone. Similar interactions between these stressors have been reported in other coral species (Nordemar et al. 2003), including early life stages (Humanes et al. 2016). Field and laboratory experiments indicate that elevated nutrients increase coral susceptibility to heat stress (Wiedenmann et al. 2013; D’Angelo et al. 2014; Burkepile et al. 2019), but the mechanisms involved are still debated. Previous work found that phosphate starvation under excess nitrogen promotes the replacement of phospholipids in the thylakoid membranes by sulfolipids, increasing susceptibility to heat and light stress in the algal symbionts (Wiedenmann et al. 2013). However, in our study, the addition of PO_4_ (NH_4_ + PO_4_ treatment) did not improve coral performance compared with corals in elevated NH_4_ alone. Alternatively, higher symbiont densities under elevated nutrients could result in a CO_2_ shortage, which might limit the dark reactions of photosynthesis (Wooldridge 2009; Wooldridge 2013), and increase algal sensitivity to photodamage (Jones et al. 1998). More research on the effects of different nutrient sources and concentrations on the nutritional state of the coral holobiont may help elucidate the mechanisms by which nutrient pollution reduces coral resistance to heat.

Heat stress has been noted as the dominant driver of prokaryotic beta-diversity change in corals facing multiple stressors (McDevitt-Irwin et al. 2019). In our study, heat stress had a large impact on the microbial community, but this stressor resulted in distinct beta-diversity changes among the corals pre-exposed to the different nutrient treatments (Fig. 5d). It is possible that these differential changes in the prokaryotic communities were associated with the increasingly unhealthy state of the corals pre-exposed to nutrients. Most of the fragments exposed to high nutrients and heat stress experienced mortality one or two days after sampling and it is possible that their prokaryotic communities were reflecting microbial activity triggered by increasing apoptosis and cell death (Yuan et al. 2017).

### Genotypes with higher performance under nutrient pollution and heat stress

Overall, genotypes *G48* and *G62*, followed by *G31*, were the most resistant to elevated nutrients alone and in combination with heat stress, exhibiting less mortality and the lowest declines in growth rates. Allocating these resistant genotypes to locations exposed to elevated nutrients might increase restoration success by reducing coral mortality directly associated with nutrient inputs and the time in which fragments would reach sizes with lower mortality risk (Goergen and Gilliam 2018). However, to achieve greater overall population resilience additional traits such as reproductive output or disease resistance should be tested to determine if there are tradeoffs between nutrient resistance and other desired characteristics (Shore-Maggio et al. 2018).

In this study, nutrient resistant genotypes were characterized by less diverse microbial communities (Fig. 5c) highly dominated by the family *Midichloriaceae* (ASV 2095; phylum Proteobacteria and genus MD3-55), and by the presence of *Spirochaetaceae* (ASV 825; phylum Spirochaetes and genus *Spirochaeta*; Fig. 5b). This contrasts with previous findings which suggest that *Midichloriaceae* is correlated with reduced growth rates under nutrient stress (Shaver et al., 2017) and is parasitic in nature because it consumes nutrients from *A. cervicornis* (Klinges 2019). We hypothesize that these discrepancies could be related to specific *Midichloriaceae* strains that associate with different *A. cervicornis* genotypes, which may respond differently to elevated nutrients. Different strains of *Midichloriaceae* in *A. cervicornis* are found across the Caribbean and the higher positive selection (or mutation rates) of this bacterium in Florida may have led to strains that associate with specific genotypes and may ultimately affect a coral’s phenotype (Baker et al. 2021). In genotypes with abundant *Midichloriaceae*, these bacteria did not increase in abundance under elevated nutrients and sometimes even decreased (Figs. 3b, 5-6). These data suggest that *Midichloriaceae* strains could play different health roles in *A. cervicornis.* This could be the case in coral genotypes that host *Midichloriaceae* at low abundances, but that experience an increase under higher nutrient availability, and compromise the coral host performance (e.g., *G07* this study, Shaver et al., 2017). In fact, the genome from the dominant *Midichloriaceae* ASV from this study has been characterized as *Candidatus* Aquarickettsia rohweri, and was sequenced from a coral with low baseline *Ca.* A. rohweri. Future studies should evaluate if *Ca*. A. rohweri that show high baseline abundances in *A. cervicornis* differ in genetic structure from those that associate with *A. cervicornis* at low abundances in Florida. Here, we suggest that *A. cervicornis* genotypes with low abundances of *Ca*. A. rohweri have a lower survivorship rate in elevated nutrients and those with high abundances are more resistant to nutrient stress and subsequent heat stress (Fig 6a).

Interestingly, *A. cervicornis* with high baseline abundances of *Midichloriaceae* have also been associated with increased disease susceptibility (Klinges 2020). Since high *Midichloriaceae* decreases disease resistance but increases survivorship under elevated nutrients, baseline *Midichloriaceae* abundances may be a biomarker for multiple stressors that could help coral practitioners make science-based outplanting decisions. As such, *A. cervicornis* with low baseline abundances of *Midichloriaceae*, like *G07* and *G08*, may be more appropriate to outplant in areas with lower nutrients and higher disease incidence, such as offshore reefs. In contrast, *A. cervicornis* with high baseline abundances of *Midichloriaceae*, like *G48, G62*, and *G31*, might be more appropriate to outplant on reefs that experience higher levels of nutrients and lower disease incidences, such as inshore reefs (Szmant and Forrester 1996; Rippe et al. 2019).

In addition to *Midichloriaceae, Spirochaetaceae* may also be used as a biomarker for survivorship in *A. cervicornis* under elevated nutrient conditions*. Spirochaetaceae* has been found in the past to distinguish *A. cervicornis* genotypes (Rosales et al. 2019). However, the implementation of either *Midichloriaceae* and *Spirochaetaceae* as biomarkers requires additional lab and field studies that test more genotypes and replicates (Parkinson et al. 2020).

## Conclusion

We found genotypic variation in the response of *A. cervicornis* to elevated nutrients, both alone and in combination with heat stress. Since *A. cervicornis* population recovery may depend heavily on the growth and survivorship rates of outplants, we suggest allocating these resistance genotypes at sites characterized by poor water quality and high nutrient loading. We also found that the prokaryotic community of *A. cervicornis* may be an indicator of nutrient resistance. While past work has suggested that *Midichloriaceae* increases with elevated nutrients, we hypothesize that only *A. cervicornis* genotypes with low baseline *Midichloriaceae* abundances increase with elevated nutrients and are more susceptible to nutrient pollution. This characteristic may be developed to screen genotypic performance of large numbers of genotypes and help select outplants based on the likelihood of survival under given environmental conditions.

## Supporting information

Electronic Supplemental Material (ESM)

