## Supplementary material for "Microbiome signatures in *Acropora cervicornis* are associated with genotypic resistance to elevated nutrients and heat stress": Electronic Supplemental Material (ESM)

### Electronic Supplementary Material for:


| | |
|---|---|
| **Supplementary methods** | **1** |
| Coral collection | 1 |
| Nutrient concentrations | 1 |
| Coral performance | 1 |
| Prokaryotic differential abundance | 2 |
| **Supplementary results** | **3** |
| Figure S1 | 3 |
| Figure S2 | 4 |
| Figure S3 | 5 |
| Figure S4 | 6 |
| Figure S5 | 7 |
| **Supplementary Tables:** | **8** |
| Table S1 | 8 |
| Table S2 | 8 |
| Table S3 | 9 |
| Table S4 | 10 |
| Table S5 | 11 |
| Table S6 | 13 |
| Table S7 | 14 |
| **Cited literature** | 16 |

### 1. Supplementary methods

#### Coral collection:

Coral fragments from six genotypes of *A. cervicornis* were donated to the University of Miami (UM) coral nursery by Mote Marine Laboratory in Summerland Key, FL, in July 2017. Replicate single-branched fragments (~ 4 cm in length) were then transported to the Marine Technology and Life Science Seawater (MTLSS) complex at the University of Miami Rosenstiel School in September 2017. Each coral was acclimated to tank conditions (26.2 ºC ± 0.6 with a 12:12 h light:dark cycle) for ~4 months. Within the first month, some fragments experienced rapid tissue loss (RTL) over a 1-2 day period, but no further mortality was observed during the following three months of acclimation. However, this initial mortality resulted in an unbalanced number of fragments tested from each genotype (N=120 fragments, 8-29 fragments per genotype; Table S1).

#### Nutrient concentrations:

Discrete water samples were collected from each nutrient treatment to verify if the addition of nutrients increased the concentrations to the target values. The samples were collected directly from the tanks using sterile 50 mL falcon tubes, immediately preserved with 100 µL of chloroform, and refrigerated at -20 ºC until they were sent for nutrient analysis. Twenty samples were analyzed for the ambient treatment and 10 for the $NH_4$ and $NH_4$ + $PO_4$ treatments.

#### Coral performance:

**Survivorship probabilities and risk of death:** Survivorship analyses were performed with the survival 2.38 (Therneau 2015) and survminer 0.4.6 (Kassambara 2018) packages for R. Survival probabilities were calculated with the Kaplan-Meier estimate (Kaplan and Meier 1958)he survival probability in a given genotype and treatment is estimated as the number of fragments that are alive in that genotype and treatment, divided by the number of fragments that were alive just prior to that time. This calculation excludes fragments that have been removed from the tanks for other studies, which are treated as "censored events". The censored events account for the incomplete information about the survivorship outcome of the removed fragments (they could have died or survived if they were left in the tanks), so the model uses the information of the "censored" individuals until the point when they are removed, but they are not considered as part of the sample groups after that day. Significant differences among the survival curves from each genotype and treatment were assessed with log-rank tests. None of the genotypes experienced mortality in the ambient treatment at any temperature phase, therefore comparisons among genotypes in the ambient treatment were not applicable. Since there were no differences between the survivorship probabilities between the $NH_4$, and $NH_4$ + $PO_4$ treatments at any temperature phase (Fig. S1), survivorship data from $NH_4$ and $NH_4$ + $PO_4$ were pooled to test for differences among the six *A. cervicornis* genotypes when exposed to elevated nutrients and further heat stress.

**Growth rates:** Growth rates were estimated using the buoyant weight technique (Davies 1989). Buoyant weight data were transformed to air weight with the formula air weight = 1/(1-water density/coral density) following (Jokiel et al. 1978). Growth rates (mg $g^{-1}$ $d^{-1}$) were estimated by calculating the difference between the air weight from two consecutive data points (mg), and normalizing this value by the initial weight of the fragment for that interval (g) and by the number of days between the two measurements (d) following previously defined methods (Ezzat et al. 2016). We first tested for the overall effect of nutrients (Ambient, $NH_4$ and $NH_4$ + $PO_4$) on *A. cervicornis* growth with a mixed-effects model that included *nutrient* treatment and number of *days* in the experiment as interacting fixed factors, and genotype and fragments as random effects (Tables S3-S4). Since there were no differences between the $NH_4$ and the $NH_4$ + $PO_4$ we pooled these treatments and tested for genotypic differences in ambient and elevated nutrients with a mixed-effects model that included *genotype*, *day*, and *nutrient* treatment (ambient versus elevated nutrients) as fixed factors, as well as coral *fragment* and *replicate* tank as a random factors (Tables S3, S5).

**Photochemical efficiency ($F_v/F_m$):** $F_v/F_m$ values were used as a proxy for the algal community function. Declining $F_v/F_m$ indicates dysfunction of the photosystem II, which can be used as an early sign of heat stress that could lead to coral bleaching (Warner, Fitt, and Schmidt 1999). Overall changes in $F_v/F_m$ associated with the nutrient treatments (ambient, $NH_4$, and $NH_4$+$PO_4$) were analyzed with a mixed-effects model that included nutrient *treatment* and number of *days* in the experiment as interacting fixed factors, as well as *genotype*, *fragments* and *replicate* tank as random effects (Table S6). Then ambient values, as well as pooled elevated nutrient values ($NH_4$ and $NH_4$+$PO_4$ treatments), were used to test for genotypic differences in $F_v/F_m$ among the six genotypes. This mixed-effects model included *genotype*, *day*, and *nutrient* treatment (ambient versus elevated nutrients) as fixed factors, and *fragments* and *replicate* tank as random effects (Table S6, S7).

#### Prokaryotic differential abundance:

##### *High-throughput 16S rRNA amplicon sequencing and bioinformatic analysis*

Small tissue samples (~3 polyps per fragment) were collected at the end of phase 1 (day 75), and during phase 3 (days 100 and 111) to characterize the microbial communities (Fig. 1). A subset of samples from each day, genotype, and nutrient treatment (N=180, Table S2) were preserved and extracted using standard organic DNA extraction protocols (Baker and Cunning 2016)16S rRNA gene V4 was amplified and sequenced using previously published primers (Apprill et al. 2015). Briefly, the samples were amplified in a 50 µL reaction using the Platinum Hot Start PCR Master Mix (2X) (ThermoFisher Scientific, Waltham, MA), 2 µL of DNA, and 1 µL of each primer with PCR run at: 1 cycle x 3 min at 94°C, 35 cycles x (45 s at 94°C, 60 s at 50°C, 90 s at 72°C), 1 cycle x 10 min at 72°C. Each PCR product was cleaned with AMPure XP beads (Beckman Coulter, Brea, CA), quantified using a Qubit™ dsDNA HS Assay Kit (TermoFisher Scientific, Waltham, MA), and normalized to 4 nM. Then 5 µL of each sample was combined into a single 1.5 mL tube. The concentration of the pool was quantified with a Qubit™ dsDNA HS Assay Kit and was below the nM threshold and thus was concentrated by using a vacuum centrifuge. The pooled sample was submitted to the Hussman Institute for Human Genomics University of Miami Miller School of Medicine and sequenced on a MiSeq with the PE-300v3 kit. Post-sequencing the data were demultiplexed at the core facility.

Demultiplexed sequences were processed on Qiime2-2018.11 (Bolyen et al. 2019). Upon initial inspection of the sequences, the reverse reads were of poor quality and thus only the forward reads were analyzed. Primers were trimmed from the forward reads with the Cutadapt plugin (Martin 2011) and then processed with the DADA2 plugin (Callahan et al. 2016). The default settings were used for DADA2 and the sequences were trimmed at the 20 and 220 bp positions. The DADA2 program output quality filtered, chimera removed Amplicon Sequence Variants (ASVs). These ASVs were taxonomically classified with a fitted classifier using the function feature-classifier classify-sklearn and a trained Silva-132-99-105-806 database (Bokulich et al. 2018). The sequences that were taxonomically assigned as chloroplast or mitochondria were removed from the analysis. An *Escherichia coli* (ASV) was found across all samples and it was the second most frequent sequence in the dataset. This ASV was removed from the analysis since it was likely a contaminant from the DNA extraction protocol which contained tRNA isolated from *E.coli* (per Millipore Sigma, Burlington, MA). Samples with <100 sequences were removed for downstream analyses (n = 1).

#### 2. Supplementary results

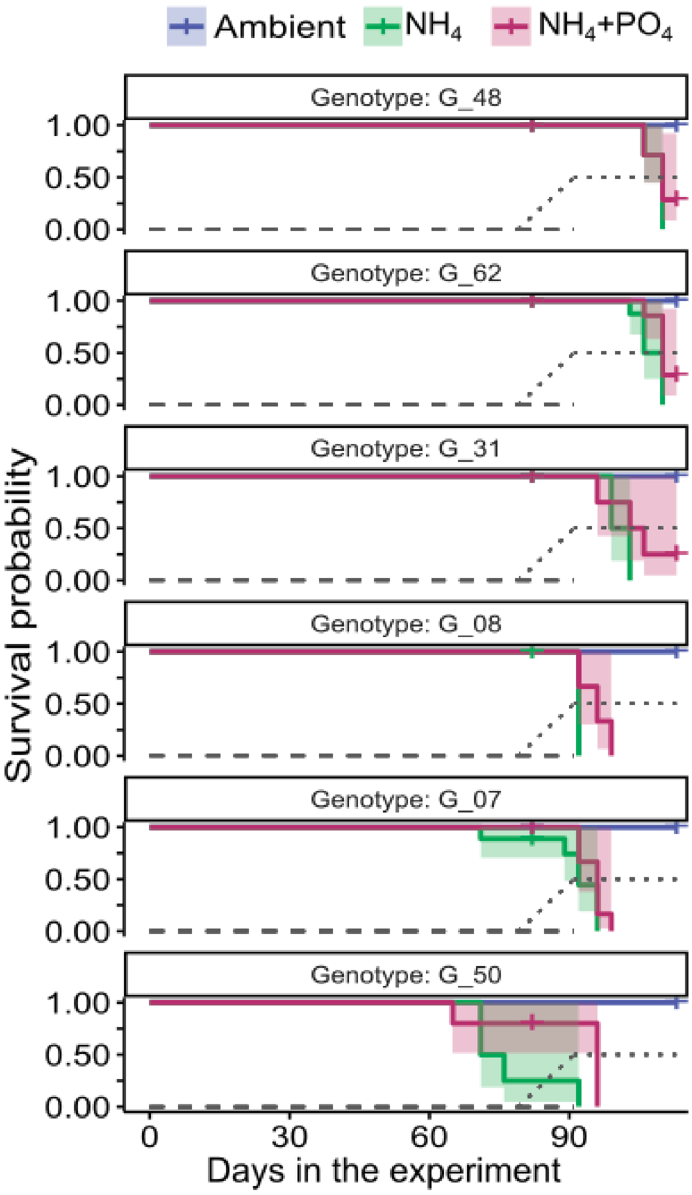


**Figure S1:** Survival probability in *A. cervicornis* exposed to nutrient treatments (ambient, $NH_4$, and $NH_4$+$PO_4$, followed by heat stress. Survival probabilities were lower for corals exposed to elevated nutrients compared to ambient nutrients (Log-rank $p < 0.0001$), but there were no significant differences between elevated $NH_4$ and $NH_4$+$PO_4$ (Log-rank $p = 0.097$). The gray dashed line represents the period of nutrient addition. The pointed lines represent the ramping up and heat stress phases.

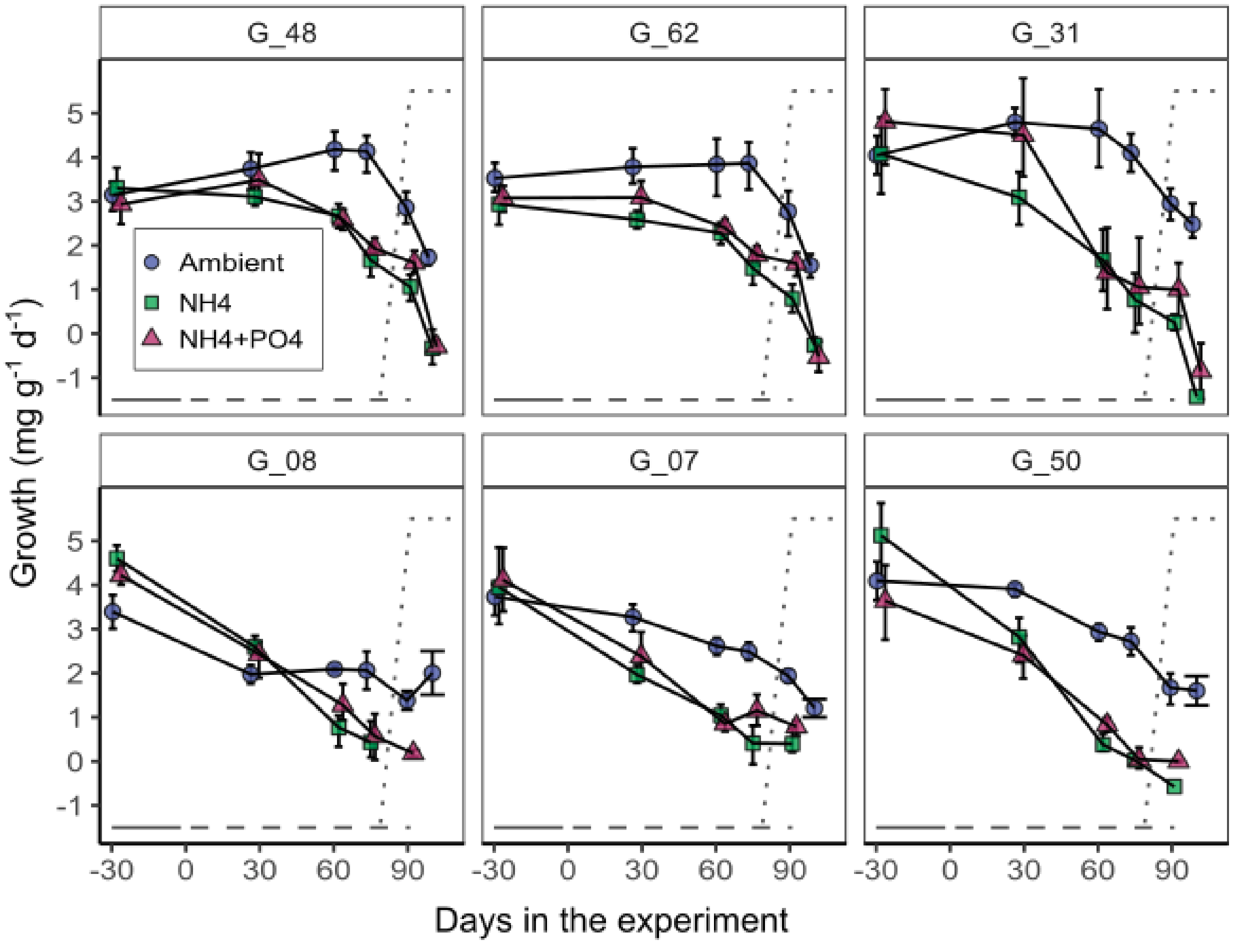


**Figure S2:** *A. cervicornis* growth rates (mean mg $g^{-1}$ $d^{-1}$ ± 95% CI) before starting the experiment (baseline), under nutrient treatments and control temperature (days 1-75), and subsequent ramp-up (days 76-90) and heat stress (days 91-113). Each panel represents a single genotype. The gray dashed line represents the period of nutrient addition. The pointed lines represent the ramping up and heat stress phases.

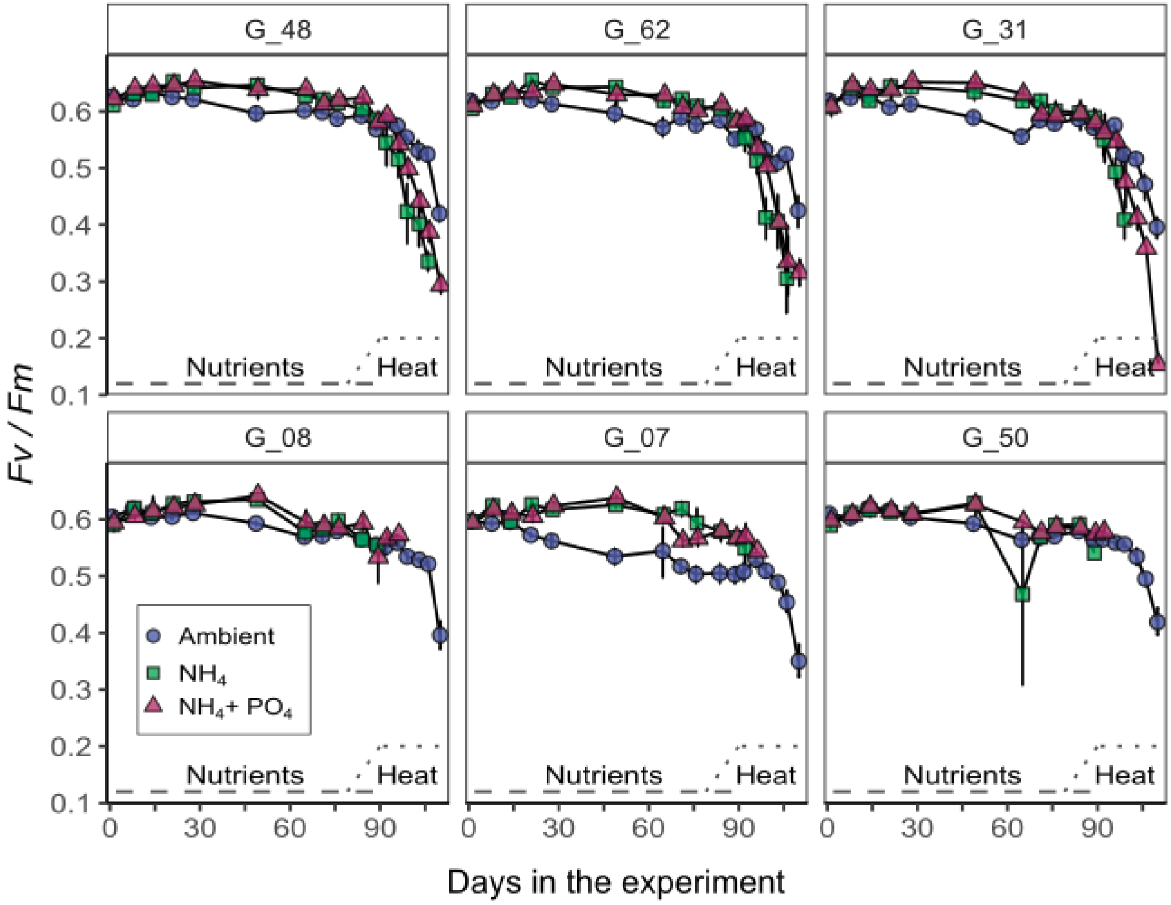


**Figure S3:** *A. cervicornis* photochemical efficiency rates (*Fv/Fm* ± 95% CI) under nutrient treatments and control temperature (days 1-75), and subsequent ramp-up (days 76-90) and heat stress (days 91-113). Each panel represents a single genotype. The gray dashed line represents the period of nutrient addition. The pointed lines represent the ramping up and heat stress phases.

***Prokaryotic differential abundance by nutrient treatment and days***

Differential abundance analysis yielded four significant ASVs (8818, 989, 2609, and 3479) among ambient, $NH_4$, and $NH_4$+$PO_4$ at control temperature (phase 1; day 75), but some of these differences were driven by specific genotypes. For ASV8818 (phylum Proteobacteria and genus *Pseudoalteromonas*) the mean RA was highest in $NH_4$+$PO_4$ (0.8% ± 1.3%), but *G31* had a particularly higher mean RA (1.9% ± 2.2%) of *Pseudoalteromonas* compared to the other genotypes. ASV989 (phylum Cyanobacteria and genus *Rivularia*), had the highest RA in ambient treatment (1.9% ±3.0%) compared to $NH_4$ (0.15% ± 0.05%), and $NH_4$+$PO_4$ (1.2% ±1.8%) but this high abundance in ambient corals was mostly driven by *G07* (3.6% ±3.2%). ASV2609 (phylum Proteobacteria and genus *Pseudomonas*), had a similar mean RA in both ambient (1.1% ± 0.9%) and $NH_4$+$PO_4$ (1.1% ± NA) and was lower in $NH_4$ (0.7% ± 0.6%). The mean RA of ASV3479 (phylum Proteobacteria genus *Methylobacterium*) was highest in ambient nutrients (1.2% ± 1.5%), followed by $NH_4$+$PO_4$ (0.9% ± 0.9%), and $NH_4$ (0.9% ± 0.7%). While not significantly different across genotypes, *G07* had a low abundance of *Midichloriaceae* (ASV 2095; 1.0% ±0.7%) at control temperature (phase 1; day 75) , but increased in $NH_4$ (20.9% ±4%) and $NH_4$+$PO_4$ (38.5±52.0) treatments.

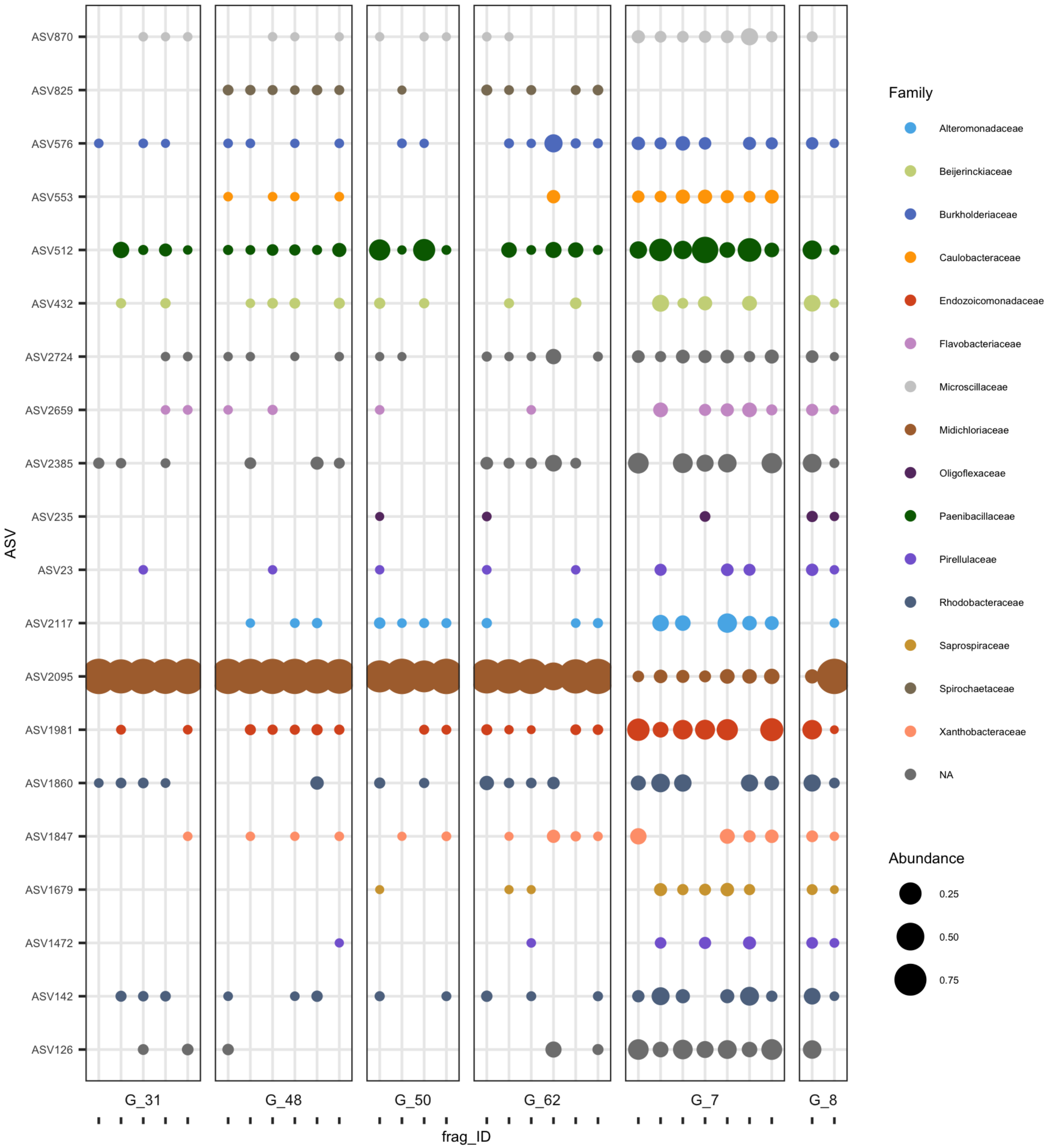


**Fig. S4: Core microbiome of six *A.cervicornis* genotypes.** The core ASVs found in at least 99% of a single genotype and then plotted across each genotype. The core members are colored by the bacteria family and the bubbles are sized based on relative abundance.

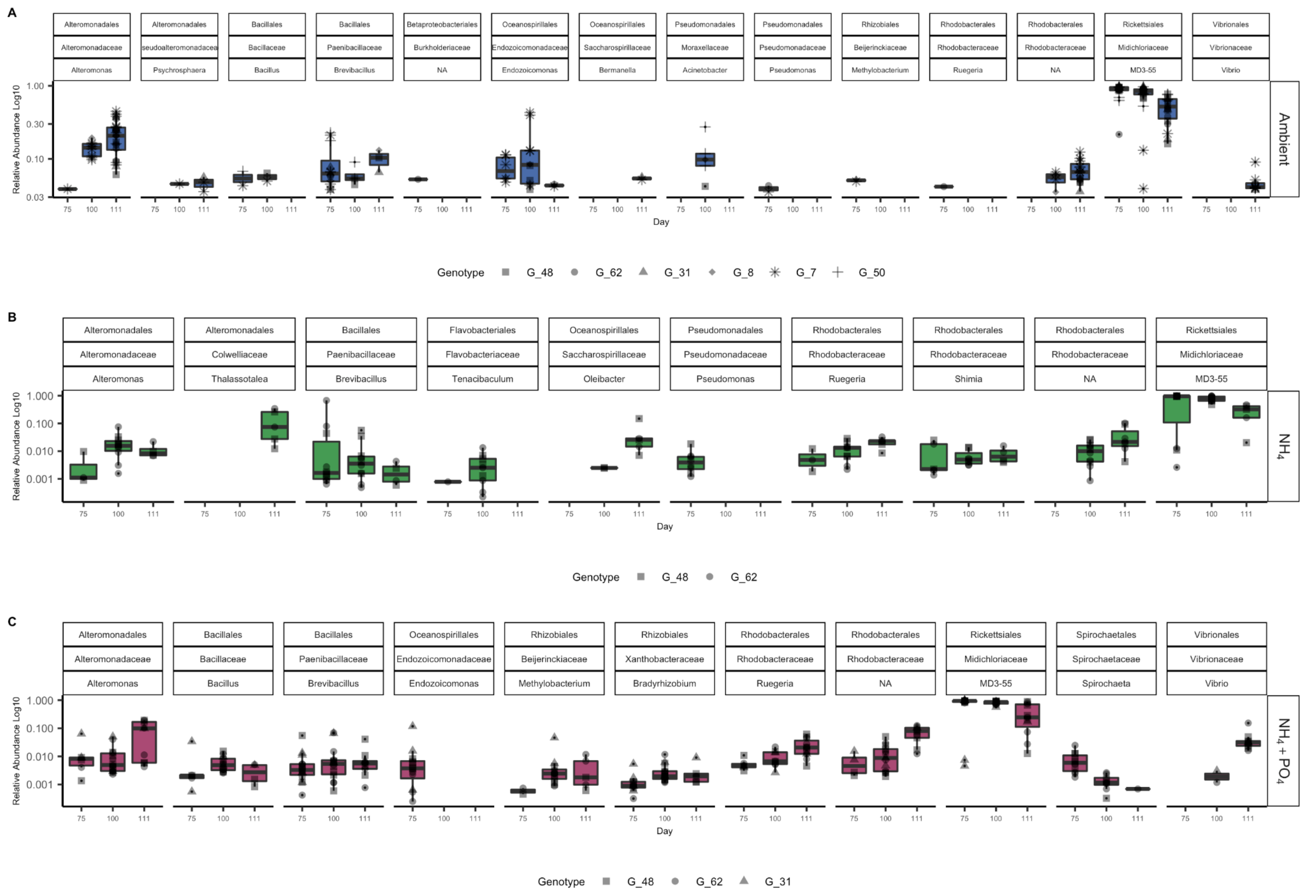


**Fig. S5:** Significantly differentiated ASVs across time and heat stress in the three nutrient treatments (A) Ambient (B) $NH_4$, and (C) $NH_4$,+ $PO_4$. Relative abundance (y-axis) of the significant taxa across time (days 75 [control], 100 [heat], and 111 [heat]). The data is parsed by the ASV's corresponding order, family, and genus. The shapes denote the six genotypes in (A), the two genotypes in (B), and the three genotypes in (C) that were used in each analysis (due to survivors across time). The genotypes are ordered by survivorship rates in each key.

### 3. Supplementary Tables:

**Table S1:** Number of fragments exposed to each nutrient treatment per *A. cervicornis* genotype. Each treatment was replicated in two independent aquaria (Tank 1 and Tank 2).

| Genotype | Treatment / Replicate tank | | | | | | Fragments per genotype |
|---|---|---|---|---|---|---|---|
| | **A** (ambient nutrients) | | **N** [ambient nutrients + 10μM $NH_4$] | | **N+P** [ambient nutrients + 10μM $NH_4$ + 1μM $PO_4$] | | |
| | **Tank 1** | **Tank 2** | **Tank 1** | **Tank 2** | **Tank 1** | **Tank 2** | |
| G48 | 5 | 5 | 5 | 4 | 5 | 4 | **28** |
| G62 | 5 | 4 | 5 | 5 | 5 | 5 | **29** |
| G31 | 3 | 2 | 3 | 3 | 3 | 2 | **16** |
| G08 | 1 | 1 | 1 | 2 | 1 | 2 | **8** |
| G07 | 5 | 4 | 5 | 4 | 4 | 4 | **26** |
| G50 | 2 | 2 | 2 | 2 | 3 | 2 | **13** |
| **Fragments per treatment** | **21** | **18** | **21** | **20** | **21** | **19** | **120** |

**Table S2:** Number of fragments per genotype used to characterize the prokaryotic communities in *A. cervicornis*. Fragments sampled in day 75 were re-sampled in days 100 and 111. Reduction in sample size overtime correspond with mortality of the fragments before they could be sampled again.

| Genotype | Day 75 | Day 100 | Day 111 | Total samples per genotype |
|---|---|---|---|---|
| G48 | 16 | 17 | 13 | **46** |
| G62 | 18 | 15 | 13 | **46** |
| G31 | 15 | 6 | 4 | **25** |
| G08 | 8 | 3 | 1 | **12** |
| G07 | 16 | 7 | 7 | **30** |
| G50 | 12 | 4 | 4 | **20** |
| **Total samples per time point** | **85** | **52** | **42** | **179** |

**Table S3**: Generalized linear mixed models used to test for differences in the growth rate of *A. cervicornis* exposed to nutrient treatments at control temperature (days 1-78) and heat stress (days 90-113). Factor "Nutrient" in Model 1 has three levels (Ambient, $NH_4$ and $NH_4$ + $PO_4$), while factor "Nutrients2" in Model 2 has two levels (Ambient, and elevated nutrients [$NH_4$ and $NH_4$ + $PO_4$ pooled]).

**Model 1: *A. cervicornis* growth with three nutrient levels**

| **Fixed effects** | **numDF** | **denDF** | **F-value** | **p-value** |
|---|---|---|---|---|
| Nutrient | 2 | 118.2 | 123.6 | <0.001 |
| Days | 5 | 491.7 | 247.3 | <0.001 |
| Nutrient:Day | 10 | 492.6 | 22.1 | <0.001 |
| **Random effects** | **npar** | **logLik** | **AIC** | **Pr(>Chisq)** |
| none | 22 | -728.7 | 1501.3 | |
| Genotype | 21 | -746.3 | 1534.6 | <0.001 |
| Fragment | 21 | -747.9 | 1537.8 | <0.001 |
| Replicate (Tank) | 21 | -728.9 | 1499.7 | 0.54 |

**Model 2: *A. cervicornis* growth with both elevated nutrient treatments pooled**

| **Fixed effects** | **numDF** | **denDF** | **F-value** | **p-value** |
|---|---|---|---|---|
| Genotype | 5 | 104.8 | 12.3 | <0.001 |
| Nutrients2 | 1 | 111.6 | 171.7 | <0.001 |
| Days | 5 | 441.2 | 283.9 | <0.001 |
| Genotype:Days | 5 | 106.9 | 3.2 | <0.01 |
| Treatment2:Days | 25 | 440.2 | 14.5 | <0.001 |
| Nutrient:Day | 5 | 441.9 | 71.8 | <0.001 |
| Genotype:Treatment2:Days | 22 | 440.4 | 2.9 | <0.001 |
| **Random effects** | **npar** | **logLik** | **AIC** | **Pr(>Chisq)** |
| none | 72 | -555.1 | 1254.4 | |
| Fragment | 71 | -622.1 | 1386.1 | <0.001 |
| Replicate (Tank) | 71 | -555.3 | 1252.7 | 0.6 |

**Table S4:** Estimated growth rates (mg $g^{-1}$ $d^{-1}$) for *A. cervicornis* exposed to nutrient treatments, and subsequent heat stress using Model 1. Pairwise comparisons between groups were obtained using Tukey's HSD test ($\alpha$ = 0.05). Stars (*) in the treatments denote the group of corals that were assigned to these treatments, but that were not exposed to elevated nutrients at the time of the measurement. Percentages of change in bold represent comparison among values that were significantly different based on the Tukey's HSD test. The model includes *nutrient* treatment, and *days* in the experiment as interacting fixed factors, as well as *genotype*, *fragment*, and *replicate* tank as random effects (see Table S3).

| Days in the experiment (Phase) | Nutrient Treatment | Em mean | SE | df | Lower CL | Upper CL | Tukey Group | % change respect ambient (same day) | % change respect baseline (day -28) | % change respect control temp (day 75) |
|---|---|---|---|---|---|---|---|---|---|---|
| -77 to -28 (Baseline) | Ambient | 3.52 | 0.2 | 11.3 | 3.05 | 3.98 | 7 | NA | NA | NA |
| | * $NH_4$ | 3.64 | 0.2 | 11.1 | 3.18 | 4.10 | 7 | 3.5% | NA | NA |
| | *$NH_4$+$PO_4$ | 3.53 | 0.2 | 11.2 | 3.07 | 3.99 | 7 | 0.4% | NA | NA |
| 1 to 28 (Control) | Ambient | 3.61 | 0.2 | 11.5 | 3.14 | 4.07 | 7 | NA | 2.6% | NA |
| | $NH_4$ | 2.59 | 0.2 | 10.9 | 2.13 | 3.05 | 6 | **-28.1%** | **-26.3%** | NA |
| | $NH_4$+$PO_4$ | 3.02 | 0.2 | 11.0 | 2.55 | 3.48 | 6-7 | -16.4% | -14.2% | NA |
| 29 to 62 (Control) | Ambient | 3.50 | 0.2 | 11.5 | 3.03 | 3.97 | 7 | NA | -0.5% | NA |
| | $NH_4$ | 1.64 | 0.2 | 10.9 | 1.18 | 2.10 | 4-5 | -53.2% | **-53.4%** | NA |
| | $NH_4$+$PO_4$ | 1.67 | 0.2 | 11.2 | 1.20 | 2.13 | 5 | -52.3% | **-52.5%** | NA |
| 62 to 75 (Control) | Ambient | 3.36 | 0.2 | 11.3 | 2.89 | 3.82 | 7 | NA | -4.4% | NA |
| | $NH_4$ | 0.96 | 0.2 | 11.7 | 0.49 | 1.42 | 2-3 | **-71.5%** | **-72.8%** | NA |
| | $NH_4$+$PO_4$ | 1.24 | 0.2 | 11.2 | 0.78 | 1.70 | 3-5 | **-63.1%** | **-64.7%** | NA |
| 75 to 91 (Ramp-up) | Ambient | 2.36 | 0.2 | 12.9 | 1.89 | 2.84 | 6 | NA | **-32.8%** | **-29.7%** |
| | $NH_4$ | 0.46 | 0.2 | 17.7 | -0.04 | 0.96 | 2 | **-80.5%** | **-86.9%** | -51.9% |
| | $NH_4$+$PO_4$ | 0.98 | 0.2 | 14.3 | 0.50 | 1.46 | 2-4 | **-58.5%** | **-72.1%** | -21.0% |
| 91 to 100 (Heat) | Ambient | 1.63 | 0.2 | 12.9 | 1.15 | 2.10 | 3-5 | NA | **-53.7%** | **-51.6%** |
| | * $NH_4$ | -0.69 | 0.3 | 26.0 | -1.23 | -0.15 | 1 | **-142.6%** | **-119.7%** | **-172.4%** |
| | *$NH_4$+$PO_4$ | -0.85 | 0.3 | 24.2 | -1.38 | -0.32 | 1 | **-152.3%** | **-124.2%** | **-168.5%** |

**Table S5:** Estimated growth rates (mg $g^{-1}$ $d^{-1}$) for six *A. cervicornis* genets exposed to nutrient treatments, and subsequent heat stress using Model 2. Pairwise comparisons between groups were obtained using Tukey's HSD test (α = 0.05). Stars (*) in the treatments denote the group of corals that were assigned to these treatments, but that were not exposed to elevated nutrients at the time of the measurement. Percentages of change in bold represent comparison among values that were significantly different based on the Tukey's HSD test. The model includes *genet*, *nutrient* treatment, and *days* in the experiment as interacting fixed factors, as well as *fragment*, and *replicate* tank as random effects (see Table S3).

| Days in the experiment (Phase) | Nutrient Treatment | Genet | Em mean | SE | df | Lower CL | Upper CL | % change respect ambient (same day) | % change respect baseline | % change respect control temp (Day 75) | % respect G_48 | % respect G_50 |
|---|---|---|---|---|---|---|---|---|---|---|---|---|
| -77 to -28 (Baseline) | Ambient | G_48 | 3.14 | 0.22 | 90.6 | 2.71 | 3.58 | NA | NA | NA | NA | -23.2% |
| | | G_62 | 3.52 | 0.23 | 104.2 | 3.07 | 3.97 | NA | NA | NA | 12.0% | -14.0% |
| | | G_31 | 4.04 | 0.30 | 193.4 | 3.44 | 4.64 | NA | NA | NA | 28.4% | -1.4% |
| | | G_08 | 3.39 | 0.48 | 285.8 | 2.45 | 4.33 | NA | NA | NA | 7.9% | -17.1% |
| | | G_07 | 3.73 | 0.23 | 104.2 | 3.27 | 4.18 | NA | NA | NA | 18.6% | -8.9% |
| | | G_50 | 4.09 | 0.34 | 226.3 | 3.42 | 4.76 | NA | NA | NA | 30.2% | NA |
| | Nutrients (N and N+P pooled) | G_48 | 3.11 | 0.17 | 36.1 | 2.78 | 3.45 | -1.0% | NA | NA | NA | -27.5% |
| | | G_62 | 3.01 | 0.16 | 30.4 | 2.68 | 3.33 | -14.6% | NA | NA | -3.5% | -30.0% |
| | | G_31 | 4.41 | 0.21 | 78.8 | 3.99 | 4.82 | 9.1% | NA | NA | 41.5% | 2.6% |
| | | G_08 | 4.38 | 0.32 | 239.9 | 3.74 | 5.02 | 29.1% | NA | NA | 40.6% | 2.0% |
| | | G_07 | 4.02 | 0.17 | 39.8 | 3.68 | 4.37 | 7.9% | NA | NA | 29.3% | -6.3% |
| | | G_50 | 4.29 | 0.23 | 104.2 | 3.84 | 4.75 | 4.9% | NA | NA | 37.9% | NA |
| 1 to 28 (Control) | Ambient | G_48 | 3.74 | 0.22 | 90.6 | 3.30 | 4.17 | NA | 18.8% | NA | NA | -4.4% |
| | | G_62 | 3.64 | 0.24 | 118.6 | 3.17 | 4.12 | NA | 3.6% | NA | -2.4% | -6.7% |
| | | G_31 | 4.79 | 0.30 | 193.4 | 4.19 | 5.39 | NA | 18.6% | NA | 28.1% | 22.5% |
| | | G_08 | 1.98 | 0.48 | 285.8 | 1.04 | 2.91 | NA | **-41.8%** | NA | -47.1% | -49.5% |
| | | G_07 | 3.27 | 0.23 | 104.2 | 2.81 | 3.72 | NA | -12.3% | NA | -12.5% | -16.4% |
| | | G_50 | 3.91 | 0.34 | 226.3 | 3.24 | 4.58 | NA | -4.5% | NA | 4.6% | NA |
| | Nutrients (N and N+P pooled) | G_48 | 3.29 | 0.17 | 36.1 | 2.96 | 3.63 | -11.9% | 5.7% | NA | NA | 27.3% |
| | | G_62 | 2.84 | 0.16 | 30.4 | 2.51 | 3.16 | -22.2% | -5.7% | NA | -13.8% | 9.7% |
| | | G_31 | 3.73 | 0.21 | 78.8 | 3.31 | 4.14 | -22.1% | -15.4% | NA | 13.2% | 44.2% |
| | | G_08 | 2.52 | 0.28 | 162.5 | 1.97 | 3.08 | **27.8%** | -42.3% | NA | -23.3% | -2.3% |
| | | G_07 | 2.16 | 0.17 | 39.8 | 1.82 | 2.50 | **-33.9%** | -46.3% | NA | -34.3% | -16.4% |
| | | G_50 | 2.58 | 0.23 | 104.2 | 2.13 | 3.04 | **-33.9%** | -39.8% | NA | -21.5% | NA |
| 29 to 62 (Control) | Ambient | G_48 | 4.18 | 0.22 | 90.6 | 3.74 | 4.61 | NA | **32.8%** | NA | NA | 42.1% |
| | | G_62 | 3.83 | 0.23 | 104.2 | 3.38 | 4.29 | NA | 8.9% | NA | -8.2% | 30.5% |
| | | G_31 | 4.63 | 0.30 | 193.4 | 4.03 | 5.23 | NA | 14.8% | NA | 11.0% | 57.7% |
| | | G_08 | 2.09 | 0.48 | 285.8 | 1.15 | 3.03 | NA | **-38.4%** | NA | -49.9% | -28.8% |
| | | G_07 | 2.61 | 0.24 | 119.0 | 2.14 | 3.08 | NA | **-30.0%** | NA | -37.5% | -11.2% |
| | | G_50 | 2.94 | 0.34 | 226.3 | 2.27 | 3.61 | NA | **-28.2%** | NA | -29.6% | NA |
| | Nutrients (N and N+P pooled) | G_48 | 2.62 | 0.17 | 36.1 | 2.29 | 2.96 | **-37.2%** | -15.8% | NA | NA | 446.3% |
| | | G_62 | 2.34 | 0.16 | 30.4 | 2.02 | 2.66 | **-39.0%** | -22.2% | NA | -10.8% | 387.2% |
| | | G_31 | 1.53 | 0.21 | 78.8 | 1.12 | 1.95 | **-66.9%** | -65.2% | NA | -41.5% | 219.6% |
| | | G_08 | 1.03 | 0.28 | 162.5 | 0.48 | 1.58 | **-50.6%** | -76.4% | NA | -60.6% | 115.2% |
| | | G_07 | 0.95 | 0.17 | 39.8 | 0.61 | 1.30 | **-63.5%** | -76.3% | | -63.7% | 98.4% |
| | | G_50 | 0.48 | 0.24 | 120.3 | 0.01 | 0.95 | **-83.7%** | -88.8% | NA | -81.7% | NA |

**Table S5 (continuation):** Estimated growth rates (mg $g^{-1}$ $d^{-1}$) for six *A. cervicornis* genets exposed to nutrient treatments, and subsequent heat stress using Model 2. Stars (*) in the treatments denote the group of corals that were assigned to these treatments, but that were not exposed to elevated nutrients at the time of the measurement. Percentages of change in bold represent comparison among values that were significantly different based on the Tukey's HSD test. The model includes *genet*, *nutrient* treatment, and *days* in the experiment as interacting fixed factors, as well as *fragment*, and *replicate* tank as random effects (see Table S3).

| Days in the experiment (Phase) | Nutrient Treatment | Genet | Em mean | SE | df | Lower CL | Upper CL | % change respect ambient (same day) | % change respect baseline | % change respect control temp (Day 75) | % respect G_48 | % respect G_50 |
|---|---|---|---|---|---|---|---|---|---|---|---|---|
| 62 to 75 (Control) | Ambient | G_48 | 4.14 | 0.22 | 90.6 | 3.70 | 4.57 | NA | **31.5%** | NA | NA | **52.1%** |
| | | G_62 | 3.86 | 0.23 | 104.2 | 3.40 | 4.31 | NA | 9.6% | NA | -6.7% | **41.9%** |
| | | G_31 | 4.09 | 0.30 | 193.4 | 3.49 | 4.69 | NA | 1.4% | NA | -1.0% | **50.5%** |
| | | G_08 | 2.06 | 0.48 | 285.8 | 1.12 | 3.00 | NA | **-39.2%** | NA | **-50.1%** | -24.2% |
| | | G_07 | 2.49 | 0.23 | 104.2 | 2.03 | 2.94 | NA | **-33.3%** | NA | **-39.9%** | -8.6% |
| | | G_50 | 2.72 | 0.34 | 226.3 | 2.05 | 3.39 | NA | **-33.5%** | NA | **-34.2%** | NA |
| | Nutrients (N and N+P pooled) | G_48 | 1.79 | 0.17 | 36.1 | 1.46 | 2.13 | **-56.7%** | -42.4% | NA | NA | -1477.9% |
| | | G_62 | 1.63 | 0.16 | 30.4 | 1.31 | 1.95 | **-57.7%** | -45.7% | NA | -8.9% | -1355.7% |
| | | G_31 | 0.90 | 0.21 | 78.8 | 0.48 | 1.31 | **-78.1%** | -79.7% | NA | -50.0% | -788.9% |
| | | G_08 | 0.51 | 0.28 | 162.5 | -0.04 | 1.06 | **-75.1%** | -88.3% | NA | -71.4% | -494.7% |
| | | G_07 | 0.79 | 0.17 | 42.9 | 0.44 | 1.13 | **-68.4%** | -80.5% | NA | -56.1% | -704.4% |
| | | G_50 | -0.13 | 0.28 | 204.8 | -0.69 | 0.43 | **-104.8%** | -103.0% | NA | -107.3% | 0.0% |
| 75 to 91 (Ramp-up) | Ambient | G_48 | 2.90 | 0.23 | 117.4 | 2.43 | 3.36 | NA | -7.8% | -29.94% | NA | 73.2% |
| | | G_62 | 2.82 | 0.25 | 138.8 | 2.33 | 3.31 | NA | -19.9% | -26.95% | -2.7% | 68.5% |
| | | G_31 | 3.07 | 0.33 | 241.9 | 2.43 | 3.72 | NA | -23.9% | -24.94% | 6.0% | 83.7% |
| | | G_08 | 1.38 | 0.48 | 285.8 | 0.44 | 2.32 | NA | -59.3% | -33.01% | -52.3% | -17.4% |
| | | G_07 | 1.88 | 0.25 | 138.7 | 1.39 | 2.37 | NA | -49.6% | -24.46% | -35.2% | 12.3% |
| | | G_50 | 1.67 | 0.34 | 226.3 | 1.00 | 2.34 | NA | -59.1% | -38.50% | -42.3% | NA |
| | Nutrients (N and N+P pooled) | G_48 | 1.43 | 0.18 | 53.8 | 1.07 | 1.80 | **-50.5%** | -53.9% | -19.98% | NA | -502.2% |
| | | G_62 | 1.19 | 0.17 | 43.4 | 0.85 | 1.54 | **-57.6%** | -60.3% | -26.88% | -16.7% | -434.9% |
| | | G_31 | 0.55 | 0.23 | 115.0 | 0.09 | 1.01 | **-82.0%** | -87.5% | -38.29% | -61.4% | -255.1% |
| | | G_08 | 0.11 | 0.42 | 418.4 | -0.72 | 0.95 | **-91.9%** | -97.4% | -78.21% | -92.2% | -131.4% |
| | | G_07 | 0.43 | 0.22 | 103.3 | -0.01 | 0.87 | **-77.3%** | -89.4% | -45.74% | -70.3% | -219.6% |
| | | G_50 | -0.36 | 0.31 | 255.0 | -0.97 | 0.25 | **-121.3%** | -108.3% | 174.16% | -124.9% | NA |
| 91 to 100 (Heat) | Ambient | G_48 | 1.77 | 0.23 | 117.4 | 1.31 | 2.24 | NA | -43.6% | -57.14% | NA | 10.5% |
| | | G_62 | 1.60 | 0.25 | 138.8 | 1.11 | 2.09 | NA | -54.6% | -58.58% | -9.8% | -0.3% |
| | | G_31 | 2.60 | 0.33 | 241.9 | 1.96 | 3.25 | NA | -35.6% | -36.44% | 46.8% | 62.3% |
| | | G_08 | 2.01 | 0.48 | 285.8 | 1.07 | 2.94 | NA | -40.9% | -2.74% | 13.2% | 25.1% |
| | | G_07 | 1.15 | 0.25 | 138.7 | 0.65 | 1.64 | NA | -69.3% | -53.90% | -35.3% | -28.5% |
| | | G_50 | 1.60 | 0.34 | 226.3 | 0.94 | 2.27 | NA | -60.8% | -41.05% | -9.5% | 0.0% |
| | Nutrients (N and N+P pooled) | G_48 | -0.27 | 0.18 | 49.1 | -0.63 | 0.09 | **-115.0%** | -108.6% | -114.88% | NA | NA |
| | | G_62 | -0.38 | 0.18 | 47.3 | -0.74 | -0.03 | **-123.8%** | -112.7% | -123.32% | 42.7% | NA |
| | | G_31 | -1.70 | 0.34 | 317.9 | -2.37 | -1.03 | **-165.3%** | -138.6% | -289.62% | 536.9% | NA |
| | | G_08 | NA | NA | NA | NA | NA | NA | NA | NA | NA | NA |
| | | G_07 | NA | NA | NA | NA | NA | NA | NA | NA | NA | NA |
| | | G_50 | NA | NA | NA | NA | NA | NA | NA | NA | NA | NA |

**Table S6**: Generalized linear mixed models used to test for differences in the photochemical efficiency (*Fv/Fv*) of *A. cervicornis* exposed to nutrient treatments at control temperature (days 1-78) and heat stress (days 90-113). Factor "Nutrient" in Model 1 has three levels (Ambient, $NH_4$ and $NH_4 + PO_4$), while factor "Nutrients2" in Model 2 has two levels (Ambient, and elevated nutrients [$NH_4$ and $NH_4 + PO_4$ pooled]).

| **Model 1: *A. cervicornis Fv/Fm* with three nutrient levels** | | | | |
|---|---|---|---|---|
| **Fixed effects** | **numDF** | **denDF** | **F-value** | **p-value** |
| Nutrient | 2 | 120.5 | 19.6 | <0.001 |
| Days | 16 | 1460.5 | 666.7 | <0.001 |
| Nutrient:Day | 31 | 1460.6 | 69.3 | <0.001 |
| **Random effects** | **npar** | **logLik** | **AIC** | **Pr(>Chisq)** |
| none | 54 | 3541.1 | -6974.2 | |
| Genotype | 53 | 3541.1 | -6899.2 | <0.001 |
| Fragment | 53 | 3465.9 | -6825.8 | <0.001 |
| Replicate (Tank) | 53 | 3502.6 | -6974.8 | 0.080 |

| **Model 2: *A. cervicornis Fv* with both elevated nutrient treatments pooled** | | | | |
|---|---|---|---|---|
| **Fixed effects** | **numDF** | **denDF** | **F-value** | **p-value** |
| Genotype | 5 | 111.1 | 35.4 | <0.001 |
| Nutrients2 | 1 | 136.8 | 22.7 | <0.001 |
| Days | 16 | 1329.7 | 316.8 | <0.001 |
| Genotype:Nutrients | 5 | 116.8 | 10.6 | <0.01 |
| Genotype:Days | 80 | 1326.6 | 2.2 | <0.001 |
| Nutrient:Days | 16 | 1331.0 | 50.1 | <0.001 |
| Genotype:Nutrients2:Days | 67 | 1327.4 | 1.7 | <0.001 |
| **Random effects** | **npar** | **logLik** | **AIC** | **Pr(>Chisq)** |
| none | 194 | 3134.2 | 5880.5 | |
| Fragment | 193 | 3133.0 | 5880.0 | <0.001 |
| Replicate (Tank) | 193 | 3094.7 | 5883.3 | 0.11 |

**Table S7:** Estimated *Fv/Fm* for six *A. cervicornis* genets exposed to nutrient treatments, and subsequent heat stress using Model 2 (see Table S6). Stars (*) in the treatments denote the group of corals that were assigned to these treatments, but that were not exposed to elevated nutrients at the time of the measurement. Percentages of change in bold represent comparison among values that were significantly different based on the Tukey's HSD test. The model includes *genet*, *nutrient* treatment, and *days* in the experiment as interacting fixed factors, as well as *fragment*, and *replicate* tank as random effects.

| Days in the experiment (Phase) | Nutrient Treatment | Genet | Em mean | SE | df | Lower CL | Upper CL | % change respect ambient (same day) | % change respect day 1 | % change respect control temp (Day 76) | % respect G_48 | % respect G_50 |
|---|---|---|---|---|---|---|---|---|---|---|---|---|
| 1 (baseline) | Ambient | G_48 | 0.63 | 0.01 | 134.52 | 0.61 | 0.64 | NA | NA | NA | NA | 2.7% |
| | | G_62 | 0.62 | 0.01 | 160.13 | 0.60 | 0.63 | NA | NA | NA | -1.2% | 1.4% |
| | | G_31 | 0.62 | 0.01 | 387.81 | 0.60 | 0.64 | NA | NA | NA | -1.2% | 1.4% |
| | | G_08 | 0.60 | 0.02 | 890.40 | 0.57 | 0.64 | NA | NA | NA | -3.4% | -0.9% |
| | | G_07 | 0.60 | 0.01 | 160.13 | 0.58 | 0.61 | NA | NA | NA | -4.7% | -2.1% |
| | | G_50 | 0.61 | 0.01 | 510.83 | 0.59 | 0.63 | NA | NA | NA | -2.6% | NA |
| | Nutrients | G_48 | 0.62 | 0.01 | 49.29 | 0.60 | 0.63 | -1.5% | NA | NA | NA | 3.8% |
| | | G_62 | 0.61 | 0.01 | 41.20 | 0.60 | 0.62 | -1.7% | NA | NA | -1.5% | 2.3% |
| | | G_31 | 0.61 | 0.01 | 114.54 | 0.59 | 0.62 | -1.7% | NA | NA | -1.4% | 2.4% |
| | | G_08 | 0.59 | 0.01 | 300.03 | 0.57 | 0.61 | -1.9% | NA | NA | -3.8% | -0.1% |
| | | G_07 | 0.60 | 0.01 | 54.38 | 0.58 | 0.61 | -0.3% | NA | NA | -3.5% | 0.2% |
| | | G_50 | 0.59 | 0.01 | 160.13 | 0.58 | 0.61 | -2.6% | NA | NA | -3.7% | NA |
| 28 (Control) | Ambient | G_48 | 0.62 | 0.01 | 134.52 | 0.61 | 0.64 | NA | -0.7% | NA | NA | 2.9% |
| | | G_62 | 0.61 | 0.01 | 160.13 | 0.60 | 0.63 | NA | -0.9% | NA | -1.4% | 1.5% |
| | | G_31 | 0.61 | 0.01 | 387.81 | 0.59 | 0.63 | NA | -0.9% | NA | -1.4% | 1.5% |
| | | G_08 | 0.61 | 0.02 | 890.40 | 0.58 | 0.65 | NA | 1.2% | NA | -1.6% | 1.3% |
| | | G_07 | 0.56 | 0.01 | 160.13 | 0.55 | 0.58 | NA | -5.8% | NA | **-9.6%** | **-6.9%** |
| | | G_50 | 0.60 | 0.01 | 510.83 | 0.58 | 0.63 | NA | -1.0% | NA | -2.8% | NA |
| | Nutrients | G_48 | 0.65 | 0.01 | 49.29 | 0.64 | 0.66 | 4.2% | 5.0% | NA | NA | 6.2% |
| | | G_62 | 0.64 | 0.01 | 41.20 | 0.63 | 0.66 | 5.2% | 6.1% | NA | -0.4% | 5.7% |
| | | G_31 | 0.65 | 0.01 | 114.54 | 0.63 | 0.66 | 5.5% | 6.4% | NA | -0.1% | 6.0% |
| | | G_08 | 0.63 | 0.01 | 300.03 | 0.61 | 0.65 | 2.9% | 6.1% | NA | -2.8% | 3.2% |
| | | G_07 | 0.62 | 0.01 | 54.38 | 0.61 | 0.63 | **10.4%** | 4.2% | NA | -4.2% | 1.7% |
| | | G_50 | 0.61 | 0.01 | 160.13 | 0.59 | 0.63 | 1.0% | 2.7% | NA | -5.8% | NA |
| 65 (Control) | Ambient | G_48 | 0.60 | 0.01 | 134.52 | 0.59 | 0.62 | NA | -3.9% | NA | 0.0% | 6.7% |
| | | G_62 | 0.57 | 0.01 | 160.13 | 0.55 | 0.59 | NA | -7.5% | NA | -5.0% | 1.3% |
| | | G_31 | 0.56 | 0.01 | 387.81 | 0.53 | 0.58 | NA | **-10.2%** | NA | -7.7% | -1.5% |
| | | G_08 | 0.57 | 0.02 | 890.40 | 0.54 | 0.60 | NA | -5.8% | NA | -5.3% | 1.0% |
| | | G_07 | 0.54 | 0.01 | 160.13 | 0.53 | 0.56 | NA | -8.8% | NA | **-9.6%** | -3.5% |
| | | G_50 | 0.56 | 0.01 | 510.83 | 0.54 | 0.59 | NA | -7.5% | NA | -6.2% | 0.0% |
| | Nutrients | G_48 | 0.63 | 0.01 | 49.29 | 0.62 | 0.65 | 5.2% | 2.7% | NA | 0.0% | 19.4% |
| | | G_62 | 0.62 | 0.01 | 41.20 | 0.61 | 0.64 | 9.1% | 2.6% | NA | -1.5% | 17.6% |
| | | G_31 | 0.62 | 0.01 | 114.54 | 0.61 | 0.64 | **12.3%** | 2.6% | NA | -1.5% | 17.6% |
| | | G_08 | 0.59 | 0.01 | 300.03 | 0.57 | 0.61 | 3.1% | -1.0% | NA | -7.3% | 10.7% |
| | | G_07 | 0.61 | 0.01 | 54.38 | 0.59 | 0.62 | **11.3%** | 1.8% | NA | -4.3% | 14.2% |
| | | G_50 | 0.53 | 0.01 | 192.86 | 0.51 | 0.55 | -6.0% | **-10.7%** | NA | **-16.2%** | 0.0% |
| 76 (Control) | Ambient | G_48 | 0.59 | 0.01 | 134.52 | 0.57 | 0.60 | NA | -6.2% | NA | 0.0% | 2.9% |
| | | G_62 | 0.58 | 0.01 | 160.13 | 0.56 | 0.59 | NA | -6.9% | NA | -2.0% | 0.9% |
| | | G_31 | 0.58 | 0.01 | 387.81 | 0.56 | 0.60 | NA | -6.5% | NA | -1.5% | 1.4% |
| | | G_08 | 0.58 | 0.02 | 890.40 | 0.55 | 0.61 | NA | -4.1% | NA | -1.3% | 1.6% |
| | | G_07 | 0.50 | 0.01 | 160.13 | 0.49 | 0.52 | NA | **-15.7%** | NA | **-14.3%** | **-11.8%** |
| | | G_50 | 0.57 | 0.01 | 510.83 | 0.55 | 0.59 | NA | -6.4% | NA | -2.9% | 0.0% |
| | Nutrients | G_48 | 0.62 | 0.01 | 49.29 | 0.61 | 0.63 | 5.2% | 0.1% | NA | 0.0% | 6.6% |
| | | G_62 | 0.61 | 0.01 | 41.20 | 0.59 | 0.62 | 5.1% | -0.4% | NA | -2.0% | 4.4% |
| | | G_31 | 0.60 | 0.01 | 114.54 | 0.58 | 0.61 | 3.3% | -1.8% | NA | -3.3% | 3.0% |
| | | G_08 | 0.59 | 0.01 | 300.03 | 0.57 | 0.61 | 2.2% | -0.2% | NA | -4.1% | 2.2% |
| | | G_07 | 0.58 | 0.01 | 59.96 | 0.57 | 0.59 | **15.3%** | -2.6% | NA | -6.1% | 0.1% |
| | | G_50 | 0.58 | 0.01 | 387.18 | 0.56 | 0.60 | 1.6% | -2.4% | NA | -6.2% | 0.0% |

**Table S7 (continuation):** Estimated *Fv/Fm* for six *A. cervicornis* genets exposed to nutrient treatments, and subsequent heat stress using Model 2 (see Table S6).

| Days in the experiment (Phase) | Nutrient Treatment | Genet | Em mean | SE | df | Lower CL | Upper CL | % change respect ambient (same day) | % change respect day 1 | % change respect control (Day 76) | % respect G_48 | % respect G_50 |
|---|---|---|---|---|---|---|---|---|---|---|---|---|
| 89 (ramp-up) | Ambient | G_48 | 0.57 | 0.01 | 191.22 | 0.55 | 0.59 | NA | -9.0% | -3.05% | 0.0% | 1.1% |
| | | G_62 | 0.55 | 0.01 | 236.28 | 0.53 | 0.57 | NA | **-10.8%** | -4.21% | -3.1% | -2.0% |
| | | G_31 | 0.57 | 0.01 | 517.23 | 0.55 | 0.59 | NA | -7.8% | -1.42% | 0.1% | 1.2% |
| | | G_08 | 0.56 | 0.02 | 890.40 | 0.52 | 0.59 | NA | -8.1% | -4.14% | -2.4% | -1.3% |
| | | G_07 | 0.50 | 0.01 | 236.28 | 0.48 | 0.52 | NA | **-15.7%** | 0.04% | **-11.6%** | **-10.6%** |
| | | G_50 | 0.56 | 0.01 | 510.83 | 0.54 | 0.59 | NA | -7.7% | -1.31% | -1.1% | 0.0% |
| | Nutrients | G_48 | 0.58 | 0.01 | 73.93 | 0.57 | 0.60 | 2.5% | -5.4% | -5.53% | 0.0% | 4.1% |
| | | G_62 | 0.58 | 0.01 | 65.45 | 0.57 | 0.60 | 6.0% | -3.8% | -3.41% | 0.2% | 4.3% |
| | | G_31 | 0.59 | 0.01 | 190.00 | 0.57 | 0.60 | 2.9% | -3.6% | -1.79% | 0.5% | 4.7% |
| | | G_08 | 0.54 | 0.01 | 387.95 | 0.52 | 0.56 | -2.3% | -8.5% | -8.35% | -7.0% | -3.2% |
| | | G_07 | 0.57 | 0.01 | 110.30 | 0.55 | 0.58 | **12.8%** | -4.6% | -2.11% | -2.7% | 1.3% |
| | | G_50 | 0.56 | 0.01 | 516.89 | 0.54 | 0.58 | -0.5% | -5.7% | -3.32% | -4.0% | 0.0% |
| 96 (Heat) | Ambient | G_48 | 0.58 | 0.01 | 191.22 | 0.56 | 0.59 | NA | -8.1% | -2.03% | 0.0% | 3.0% |
| | | G_62 | 0.57 | 0.01 | 236.28 | 0.55 | 0.59 | NA | -8.0% | -1.15% | -1.1% | 1.8% |
| | | G_31 | 0.57 | 0.01 | 517.23 | 0.55 | 0.60 | NA | -7.2% | -0.73% | -0.2% | 2.7% |
| | | G_08 | 0.56 | 0.02 | 890.40 | 0.52 | 0.59 | NA | -7.6% | -3.62% | -2.9% | 0.0% |
| | | G_07 | 0.53 | 0.01 | 236.28 | 0.51 | 0.55 | NA | **-11.3%** | 5.26% | **-8.0%** | -5.2% |
| | | G_50 | 0.56 | 0.01 | 510.83 | 0.53 | 0.58 | NA | -8.4% | -2.06% | -2.9% | 0.0% |
| | Nutrients | G_48 | 0.53 | 0.01 | 73.93 | 0.52 | 0.54 | -8.0% | -14.2% | **-14.29%** | 0.0% | NA |
| | | G_62 | 0.52 | 0.01 | 65.45 | 0.51 | 0.54 | -7.9% | -13.8% | **-13.43%** | -1.0% | NA |
| | | G_31 | 0.52 | 0.01 | 233.53 | 0.50 | 0.53 | **-10.1%** | -15.1% | **-13.57%** | -2.5% | NA |
| | | G_08 | 0.57 | 0.02 | 1282.44 | 0.53 | 0.62 | 2.4% | -3.6% | -3.43% | 8.0% | NA |
| | | G_07 | 0.54 | 0.02 | 1262.71 | 0.49 | 0.59 | 2.1% | -9.2% | -6.81% | 2.1% | NA |
| 99 (Heat) | Ambient | G_48 | 0.55 | 0.01 | 191.22 | 0.54 | 0.57 | NA | -11.4% | -5.54% | 0.0% | -0.2% |
| | | G_62 | 0.53 | 0.01 | 236.28 | 0.51 | 0.55 | NA | -13.9% | -7.51% | -4.0% | -4.2% |
| | | G_31 | 0.52 | 0.01 | 517.23 | 0.50 | 0.55 | NA | -15.5% | -9.60% | -5.8% | -6.0% |
| | | G_08 | 0.53 | 0.02 | 890.40 | 0.50 | 0.57 | NA | -11.6% | -7.77% | -3.7% | -3.9% |
| | | G_07 | 0.51 | 0.01 | 236.28 | 0.49 | 0.53 | NA | -14.6% | 1.34% | **-8.1%** | **-8.3%** |
| | | G_50 | 0.56 | 0.01 | 510.83 | 0.53 | 0.58 | NA | -8.8% | -2.54% | 0.2% | 0.0% |
| | Nutrients | G_48 | 0.46 | 0.01 | 73.93 | 0.45 | 0.47 | **-16.9%** | -25.2% | **-25.32%** | 0.0% | NA |
| | | G_62 | 0.46 | 0.01 | 65.45 | 0.44 | 0.47 | **-14.5%** | -25.1% | **-24.75%** | -1.2% | NA |
| | | G_31 | 0.45 | 0.01 | 381.88 | 0.43 | 0.47 | **-14.1%** | -26.1% | **-24.80%** | -2.6% | NA |
| 106 (Heat) | Ambient | G_48 | 0.53 | 0.01 | 191.22 | 0.51 | 0.54 | NA | -16.1% | **-10.57%** | 0.0% | 6.1% |
| | | G_62 | 0.52 | 0.01 | 236.28 | 0.51 | 0.54 | NA | -15.1% | **-8.87%** | -0.1% | 5.9% |
| | | G_31 | 0.47 | 0.01 | 517.23 | 0.45 | 0.49 | NA | **-24.0%** | **-18.76%** | **-10.6%** | -5.2% |
| | | G_08 | 0.52 | 0.02 | 890.40 | 0.49 | 0.56 | NA | -13.7% | **-10.01%** | -0.7% | 5.3% |
| | | G_07 | 0.45 | 0.01 | 236.28 | 0.44 | 0.47 | NA | **-23.8%** | **-9.67%** | **-13.5%** | **-8.2%** |
| | | G_50 | 0.50 | 0.01 | 510.83 | 0.47 | 0.52 | NA | -18.8% | **-13.19%** | -5.7% | 0.0% |
| | Nutrients | G_48 | 0.36 | 0.01 | 110.46 | 0.34 | 0.37 | **-31.8%** | -42.0% | **-42.04%** | 0.0% | NA |
| | | G_62 | 0.32 | 0.01 | 126.44 | 0.31 | 0.34 | **-38.6%** | -47.0% | **-46.79%** | **-10.0%** | NA |
| | | G_31 | 0.35 | 0.02 | 1264.08 | 0.31 | 0.40 | **-24.8%** | -41.9% | **-40.85%** | -1.3% | NA |
| 110 (Heat) | Ambient | G_48 | 0.42 | 0.01 | 191.22 | 0.40 | 0.44 | NA | -32.8% | **-28.36%** | 0.0% | 0.4% |
| | | G_62 | 0.43 | 0.01 | 236.28 | 0.41 | 0.44 | NA | -31.2% | **-26.07%** | 1.2% | 1.6% |
| | | G_31 | 0.39 | 0.01 | 517.23 | 0.37 | 0.42 | NA | **-36.2%** | **-31.78%** | -6.2% | -5.8% |
| | | G_08 | 0.40 | 0.02 | 890.40 | 0.36 | 0.43 | NA | -34.5% | **-31.67%** | -5.9% | -5.5% |
| | | G_07 | 0.35 | 0.01 | 236.28 | 0.33 | 0.37 | NA | **-41.2%** | **-30.26%** | **-16.6%** | **-16.3%** |
| | | G_50 | 0.42 | 0.01 | 510.83 | 0.39 | 0.44 | NA | -31.3% | **-26.56%** | -0.4% | 0.0% |
| | Nutrients | G_48 | 0.29 | 0.02 | 957.33 | 0.25 | 0.32 | **-31.6%** | -53.4% | **-53.42%** | 0.0% | NA |
| | | G_62 | 0.31 | 0.02 | 954.51 | 0.28 | 0.35 | **-26.3%** | -48.4% | **-48.16%** | 9.1% | NA |
| | | G_31 | 0.15 | 0.02 | 1264.08 | 0.10 | 0.19 | **-62.4%** | -75.6% | **-75.18%** | **-48.5%** | NA |

#### Cited literature

Apprill A, McNally S, Parsons R, Weber L (2015) Minor revision to V4 region SSU rRNA 806R gene primer greatly increases detection of SAR11 bacterioplankton. Aquat Microb Ecol 75:129–137

Baker AC, Cunning R (2016) Bulk gDNA extraction from coral samples. Protocols io Available at: https://www protocols io/view/Bulk-gDNA-extraction-from-coral-samples-dyq7vv

Bokulich NA, Kaehler BD, Rideout JR, Dillon M, Bolyen E, Knight R, Huttley GA, Gregory Caporaso J (2018) Optimizing taxonomic classification of marker-gene amplicon sequences with QIIME 2's q2-feature-classifier plugin. Microbiome 6:90

Bolyen E, Rideout JR, Dillon MR, Bokulich NA, Abnet CC, Al-Ghalith GA, Alexander H, Alm EJ, Arumugam M, Asnicar F, Bai Y, Bisanz JE, Bittinger K, Brejnrod A, Brislawn CJ, Brown CT, Callahan BJ, Caraballo-Rodríguez AM, Chase J, Cope EK, Da Silva R, Diener C, Dorrestein PC, Douglas GM, Durall DM, Duvallet C, Edwardson CF, Ernst M, Estaki M, Fouquier J, Gauglitz JM, Gibbons SM, Gibson DL, Gonzalez A, Gorlick K, Guo J, Hillmann B, Holmes S, Holste H, Huttenhower C, Huttley GA, Janssen S, Jarmusch AK, Jiang L, Kaehler BD, Kang KB, Keefe CR, Keim P, Kelley ST, Knights D, Koester I, Kosciolek T, Kreps J, Langille MGI, Lee J, Ley R, Liu Y-X, Loftfield E, Lozupone C, Maher M, Marotz C, Martin BD, McDonald D, McIver LJ, Melnik AV, Metcalf JL, Morgan SC, Morton JT, Naimey AT, Navas-Molina JA, Nothias LF, Orchanian SB, Pearson T, Peoples SL, Petras D, Preuss ML, Pruesse E, Rasmussen LB, Rivers A, Robeson MS 2nd, Rosenthal P, Segata N, Shaffer M, Shiffer A, Sinha R, Song SJ, Spear JR, Swafford AD, Thompson LR, Torres PJ, Trinh P, Tripathi A, Turnbaugh PJ, Ul-Hasan S, van der Hooft JJJ, Vargas F, Vázquez-Baeza Y, Vogtmann E, von Hippel M, Walters W, Wan Y, Wang M, Warren J, Weber KC, Williamson CHD, Willis AD, Xu ZZ, Zaneveld JR, Zhang Y, Zhu Q, Knight R, Caporaso JG (2019) Reproducible, interactive, scalable and extensible microbiome data science using QIIME 2. Nat Biotechnol 37:852–857

Callahan BJ, McMurdie PJ, Rosen MJ, Han AW, Johnson AJA, Holmes SP (2016) DADA2: High-resolution sample inference from Illumina amplicon data. Nat Methods 13:581–583

Davies SP (1989) Short-term growth measurements of corals using an accurate buoyant weighing technique. Mar Biol 101:389–395

Ezzat L, Towle E, Irisson J-OO, Langdon C, Ferrier-pagès C (2016) The relationship between heterotrophic feeding and inorganic nutrient availability in the scleractinian coral T . reniformis under a short-term temperature increase. Limnol Oceanogr 61:89–102

Jokiel PL, Maragos JE, Franzisket L (1978) Coral growth: buoyant weight technique. Coral reefs: research methods 529–541

Kaplan EL, Meier P (1958) Nonparametric Estimation from Incomplete Observations. J Am Stat Assoc 53:457–481

Kassambara A (2018) Kosinski M. survminer: Drawing survival curves using "ggplot2," 2018. URL https://CRAN R-project org/package= survminer R package version 0.4.3

Martin M (2011) Cutadapt removes adapter sequences from high-throughput sequencing reads. EMBnet.journal 17:10–12

Therneau T (2015) A Package for Survival Analysis in S. version 2.38.
